# Joint visual-vestibular computation of head direction and reflexive eye movement

**DOI:** 10.1101/2025.02.11.637650

**Authors:** Tamas Dalmay, Luke Ewig, Botond Roska, Rava Azeredo da Silveira

## Abstract

Several cognitive maps have been identified, but what sensory signals drive them and how these are combined are not well understood. One such map, the head-direction representation, is believed to be primarily driven by vestibular motion signals in mammals. Here, we combine in vivo imaging of neuronal activity, genetic perturbation of neuronal circuits, behavioral testing, and theoretical modeling to show that the representation of head direction in mouse is driven by not only vestibular but also visual motion signals: both are essential, and the latter, originating in direction-selective retinal ganglion cell activity, dominates at low speeds. We show that, correspondingly, visual perturbations alter navigational behavior that relies on head-direction computation. Finally, we find that head-direction representation and the slow phase of reflexive eye movement are tightly correlated, and we propose a theoretical model that elucidates their emergence from coupled visual and vestibular processes. Our results suggest that the brain’s estimate of head direction is built on an oculomotor reflex pathway driven by both visual and vestibular signals.

## Introduction

Cognitive maps were conjectured by Tolman as a way to organize knowledge in the brain to support behavior^1,2^. Cognitive maps were first observed in the context of spatial behavior, in the representation of location by place cells^3,4^ and by grid cells^5^, and in the representation of head direction by head-direction cells^6,7^. These spatial maps were found in hippocampus, entorhinal cortex, thalamus, and other brain regions, and arise from neuronal processing based on sensory cues. How signals originating from different sensory systems combine in the computation of these maps is not well understood.

Navigation is central to survival. Goal-directed navigation—such as searching for food, shelters, and mates—requires a sense of direction within the environment^8^. Head-direction cells are thought to provide such a sense^6,9–11^: each head-direction cell is most active at a given “preferred” head direction; as an animal’s head turns in the horizontal plane, different sets of head-direction cells become active, thereby providing a neuronal estimate of head direction^6,12^. Neurons with this property have been identified in several species^6,13–19^, including insects, fish, birds, bats, rodents, and primates, an observation which points to their ethological relevance.

The responses of head-direction cells are determined by different factors. Landmarks that are present in the environment define a reference frame with respect to which neuronal activity encodes the head direction^7,20,21^. This anchoring of the neuronal head-direction estimate to landmarks has been extensively studied and is known to rely primarily on visual information^7,22–24^. In addition, the brain continuously receives inputs that carry information about the animal’s angular head velocity, which can be used to update the neuronal head-direction estimate from instant to instant^9,20,21^. The velocity signals used for this instant-to-instant updating of the head-direction estimate are believed to be predominantly of vestibular origin in mammals, as perturbations of the vestibular system drastically impair the head-direction system^14,25,26^. Yet, head turns generate rotational visual motion cues which may also provide velocity information to the animal via activity in visual neurons. A previous study^27^ which investigated the interaction between visual and vestibular motion cues and their effects on the head-direction representation suggested that vestibular cues have a stronger impact than visual cues, though visual motion cues may exert a modulatory effect. In contrast, other studies have reported an influence of visual motion stimulation on the head-direction representation^28,29^. Taken together, the extent to which visual motion cues contribute to updating the head-direction estimate, the way in which they might interact with vestibular signals, and their possible impact on navigation are not well understood^11^. In this work, we address this problem without the confound of visual landmark cues that might provide positional information.

Apart from their putative interactions in the computation of head direction, visual and vestibular motion cues are known to drive reflexive eye movements through the optokinetic reflex (OKR) and vestibulo-ocular reflex (VOR)^30^. These reflexive eye movements are thought to contribute to gaze-stabilization during head motion in the horizontal plane^31^. Therefore, it is possible that the instant-to-instant updating of the head-direction system and the generation of reflexive eye movements by the oculomotor system are coupled.

In this work, we aimed to understand to the interactions between visual and vestibular signals in the updating of the brain’s head-direction estimate as well as the relationship of the latter with reflexive eye movements. In order to access the brain’s head-direction estimate, we recorded from populations of head direction cells in the anterodorsal nucleus of the thalamus (AD), the brain region in which head-direction cells are proportionally most abundant^32^. We measured the changes of that estimate in response to independent manipulations of visual and vestibular motion cues in head-fixed mice, and simultaneously tracked their eye movements.

Our results show a strong influence of visual motion cues on the head-direction estimate, independent of visual landmark cues. In head-fixed mice, visual motion cues dominate over vestibular cues at low speeds and vice-versa at high speeds; yet motion cues of both modalities are required at all speeds. We further found that, across a wide range of visual-vestibular stimulation conditions, the head-direction estimate strongly correlated with the slow phase of reflexive eye movements. Based on these findings, we proposed a theoretical model for the joint computation of head direction and reflexive eye movement. In addition to recording from wild-type animals, we also recorded from animals with a targeted mutation in the *Frmd7* gene, lacking horizontal direction-selectivity in the retina and the horizontal OKR^33^. These animals were unable to make use of visual motion cues to update their head-direction estimate, in line with model expectations. Both wild-type and *FRMD7^tm^*mice displayed similar shifts in their head-direction estimates when rotated in complete darkness. In freely moving mice, visual motion cues also influence the head-direction representation and they affect navigation behavior, and both depend on horizontal direction-selectivity in the retina. Finally, we introduce a probabilistic framework for multisensory estimation of head velocity that incorporates not only visual and vestibular but also motor signals, and demonstrate that it can explain key elements of visual-vestibular influence on head-direction representation.

## Results

### Population activity in anterodorsal thalamus encodes head direction during passive motion

We recorded neuronal activity in the mouse AD at cellular resolution, using a miniature head-mounted microscope. Neurons were labeled with the genetically encoded calcium indicator GCaMP6s expressed via an adeno-associated viral vector (AAV2/1-EF1a-GCaMP6s) injected into the AD (Fig. 1a, Extended Data Fig. 1a-c). To confirm that we recorded from the AD, we sectioned the brain after each experiment and identified the brain area under the lens of the head-mounted microscope (Extended Data Fig. 1d,e). We carried out recordings in an arena consisting of a central, rotatable motorized platform that we installed to allow for vestibular stimulation, and a surrounding 360-degree cylindrical LED screen that we used for visual stimulation.

**Fig.1.**
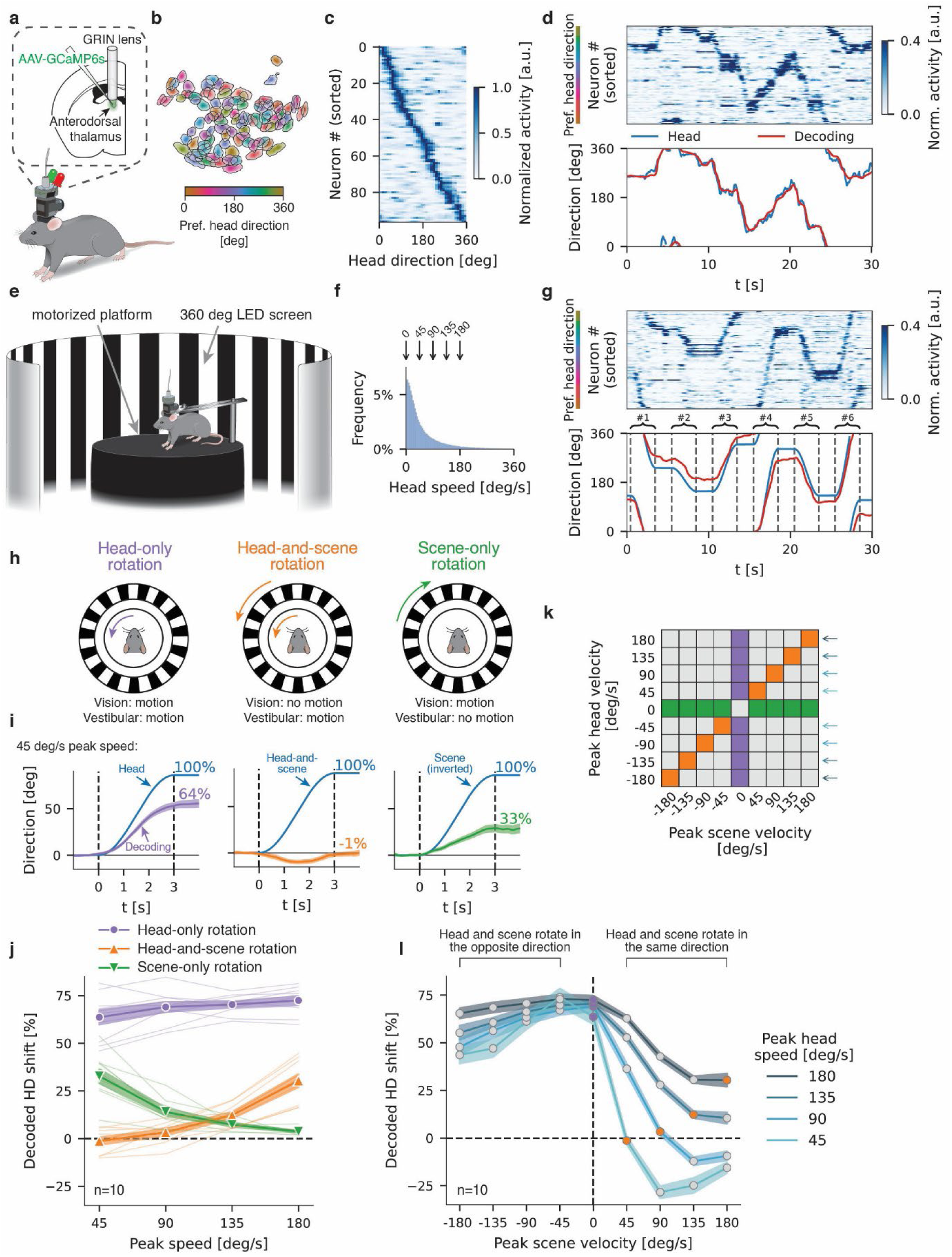
Speed-dependent control of the decoded head direction by visual and vestibular motion cues. **a**, Calcium imaging in freely moving mice with a miniature head-mounted microscope. AAV-EF1a-GCaMP6s was injected, and a GRIN lens implanted in the anterodorsal thalamus. **b**, Spatial footprints of extracted cells (n=97) from an example recording, colored according to the preferred head direction. Color intensity corresponds to the pixel weight in the footprint as quantified by CNMF-E^36,37^, the algorithm used to extract the calcium signals. **c**, Heatmap of head-direction tuning of all extracted cells from the same recording as in **b**. **d**, Top: Normalized activity of all extracted cells (sorted by preferred head direction) during a 30 second time window from the same recording as in **b**. Bottom: Actual (blue) and decoded (red) head direction for the same time window shown above. Note that the decoded head direction accurately tracks the actual head direction. **e**, Schematic of the experimental setup. Mice were head-fixed in the center of a motorized platform, surrounded by a 360° cylindrical LED screen displaying a scene consisting of a grating pattern. **f**, Histogram of angular head speeds across all mice (n=10) during freely moving recordings. Arrows indicate the peak speeds of platform rotation stimuli. **g,** Top: Normalized activity of all extracted cells (sorted by preferred head direction) during a 30 second time window from one recording in which mice were passively rotated while the scene remained stationary. Bottom: Actual (blue) and decoded (red) head direction for the same time window as shown above. **h**, Schematic of the experimental conditions for three kinds of visual and vestibular stimulation. Left: The head rotates while the scene remains stationary. Middle: Head and scene rotate in synchrony. Right: The scene rotates while the head remains stationary. **i**, Decoded head direction and stimulus angle over time for the three experimental conditions in **h**, with a peak stimulus speed of 45 deg/s. Traces show the average across animals (n=10) and the shaded area indicates the standard error of the mean. Note that data involving CW head rotations is sign-inverted (i.e., multiplied by - 1) to align with CCW head rotations, while data from CCW scene rotations is sign-inverted to align with CW scene rotations. **j**, Decoded head-direction (HD) shift as a function of the peak stimulus speed in the three experimental conditions illustrated in **h**. Thick lines and markers indicate the mean across animals (n=10), shaded areas indicate the standard error of the mean, and thin lines indicate individual mice. Decoded head-direction shift, in percent, is normalized to the angular shift of either head (purple and orange line) or scene (green line). **k**, Matrix showing the set of all experimental conditions, with purple, green and orange colors highlighting the conditions shown in **h**-**j**. **l**, Decoded head-direction shift as a function of the peak scene velocity for each non-zero value of the peak head speed. Decoded head-direction shift, in percent, is normalized to the angular head shift. Lines indicate the mean, shaded areas indicate the standard error of the mean (computed across n=10 mice). Purple and orange markers correspond to the same data as in **j**.

We first recorded neuronal activity in freely moving mice (n=10). We tracked head direction using red and green LEDs attached to the head-mounted microscope (Fig. 1a). Three geometric shapes (bar, circle, triangle) displayed on the cylindrical LED screen served as visual landmarks. 90±9% (mean across mice ± standard deviation) of the 66±27 recorded neurons exhibited head-direction tuning (Extended Data Fig. 1f), and the preferred head directions of different neurons spread uniformly over all angles (Rayleigh test: p>0.05 for each mouse) (Fig. 1b,c). The activity of AD neurons, when ordered according to their preferred head direction, took the shape of a bump that traveled in the neuronal population over time, as the mouse was turning its head while locomoting on the platform (Fig. 1d). To decode the head direction from the activity of the recorded neurons, we embedded the high-dimensional population activity in a two-dimensional space using Laplacian Eigenmaps^34^. The embedded data had a circular structure, and we defined the azimuthal coordinate (angle) of a point in the embedded data as the decoded head direction (Extended Data Fig. 1g). This was justified by the strong circular correlation (0.86±0.08) between the azimuthal coordinate of the embedded data and the head direction of the mouse. Furthermore, the azimuthal coordinate tracked the head direction (with an error of 15±6 degrees) (Fig. 1d, Extended Data Fig. 1h).

In order to examine the influence of both visual and vestibular motion cues on the decoded head direction in a controlled way, we next performed experiments in which the mouse was rotated passively. Head-fixing the mouse in the center of the motorized platform enabled us to independently control visual and vestibular motion cues by rotating the platform, the visual scene, or both. The visual scene displayed on the cylindrical LED screen consisted of a vertical black-and-white grating pattern (with a 11.25-degrees angular period, Fig. 1e). We chose such a highly symmetric pattern, free of any cue that could serve as a visual landmark, to specifically study the role of visual motion in the instant-to-instant updating of the head-direction estimate, without the confound of anchoring to visual landmarks.

We first recorded from AD while we rotated the platform but kept the scene stationary (“head-only rotation” condition). This condition is similar to the natural situation in which the visual world is stationary as a whole and global visual motion results from head motion. Each rotation lasted three seconds and had a sinusoidal velocity profile (Extended Data Fig. 1i) with a single peak. To address a behaviorally relevant range of speeds, we used five peak speeds (0 deg/s, 45 deg/s, 90 deg/s, 135 deg/s, 180 deg/s) which spanned the range most frequently exhibited by freely moving mice (Fig. 1f). This resulted in rotation angles of 86 degrees, 172 degrees, 258 degrees and 344 degrees, respectively. The clockwise (CW) and counterclockwise (CCW) rotations were presented in random order. AD activity during passive rotations was qualitatively similar to that in freely moving mouse: a bump of activity tracked the direction of the head as the platform, and hence the mouse, was rotated (Fig. 1g)^35^. The decoded head direction shifts largely matched the actual head direction shifts, though with a degree of underestimation which was possibly due to the lack of proprioceptive or motor efferent signals during passive rotations (Extended Data Fig. 1j). The access to an internal estimate of head direction in the brain, together with the ability to control visual and vestibular motion cues experimentally, provided us with a handle to investigate the effects of the two types of sensory cues on the decoded head direction.

### The decoded head direction depends on both visual and vestibular motion cues, and vision dominates at low speed

We explored vestibular and putative visual effects on the decoded head direction by independently manipulating visual and vestibular motion cues in the passive head-fixed condition. Specifically, we recorded from AD while rotating the platform or the scene, or both, according to the same velocity profiles discussed above. We initially considered three conditions, each with a different combination of platform and scene velocity profiles (Fig. 1h). In the first condition, we rotated the platform while the scene remained stationary (head-only rotation, as discussed before). Here both visual and vestibular motion cues indicate rotation. In the second condition, we rotated the platform and the scene according to the same velocity profile (“head-and-scene rotation”). Here only vestibular motion cues indicate rotation. In the third condition, we rotated the scene while the platform remained stationary (“scene-only rotation”). Here only visual motion cues indicate rotation. The last two conditions represent a conflict in that only one of the two sensory cues indicates motion while the other does not.

We observed stark differences among the decoded head directions in each of the three conditions (Fig. 1i, n=10 mice) for peak speed of 45 deg/s, which corresponds to the most frequent non-zero angular head speed (among those we tested) in freely moving mice (Fig. 1f). During head-only rotation, the decoded head direction tracked the actual head direction, with a decoded head-direction shift of 64±4% (mean across mice ± standard error of the mean) normalized to the angular shift of the head. By contrast, the decoded head direction remained mostly unchanged during head-and-scene rotation with a normalized shift of -1±2%, significantly less than the decoded head-direction shift evoked by head-only rotation (p=1.8×10^-7^, one-sided paired t-test). During scene-only rotation the decoded head direction shifted by 33±4% normalized to the angular shift of the scene (p=2.0×10^-5^, one-sided t-test for zero mean vs. positive mean). The decoded head-direction shift was sign-inverted as compared to the scene shift, consistent with the natural situation in which head motion induces global visual motion in the opposite direction to the movement of the head. These results suggest that at 45 deg/s peak speed the updating of the decoded head direction is based predominantly on visual motion cues rather than vestibular ones, as head-only and scene-only rotation elicit a much larger response in AD than head-and-scene rotation.

We asked whether this trend would be maintained at higher speed, and, more generally, how the relative impact of visual and vestibular motion cues changes as a function of speed. We compared activity in AD across all peak speeds in the three conditions (Fig. 1j, Extended Data Fig. 2a). During head-only rotation, the decoded head-direction shifts exhibited only a weak dependence on speed (0.1% per deg/s; p=0.025, two-sided t-test on the slope of a linear curve fit), with a normalized decoded head-direction shift of 69%±1% on average. During head-and-scene rotation, the normalized decoded head-direction shift was close to zero (-1±2%) at 45 deg/s peak speed, and increased (p=9.5×10^-11^) with speed up to 30±3% at 180 deg/s peak speed. Yet the decoded head-direction shift evoked by head-and-scene rotation remained significantly lower than that evoked by head-only rotation (p<10^-6^ for all speeds, one-sided paired t-test). The normalized decoded head-direction shift during scene-only rotation exhibited the opposite pattern: it decreased (p=4.6×10^-10^) with speed, from 33±4% at 45 deg/s peak speed to 4±1% at 180 deg/s peak speed. Nonetheless, it was significantly greater than zero for all speeds (p<10^-4^ for all speeds, one-sided t-test for zero mean vs. positive mean).

From these observations, two patterns emerge. First, scene-only rotations elicit greatest relative response at low speeds, at which head-and-scene rotations evoke little response; conversely, head-and-scene rotations elicit greatest relative response at high speeds, at which scene-only rotations evoke little response (Fig. 1j). This indicates that visual motion cues dominate the decoded head-direction shift at low speeds, while vestibular motion cues dominate at high speeds. Second, the decoded head-direction shift measured in head-only rotation is larger than the sum of the shifts measured in head-and-scene and scene-only rotations. Thus, visual motion cues affect the decoded head direction even at high speeds, and, conversely, vestibular motion cues affect the decoded head direction even at low speeds. The neuronal response elicited by head-only rotation, in which both the visual system and the vestibular system signal motion, cannot be obtained as a linear combination of the responses in conditions in which motion cues from one of the two modalities are abolished (Extended Data Fig. 2b).

Thus far we have examined head-direction coding in AD for particular choices of visual and vestibular stimulations, in which either one of the two velocity profiles was set to zero (head-only and scene-only rotations) or the two velocity profiles were identical (head-and-scene rotation). For a closer examination of the way in which visual and vestibular motion cues together update the decoded head direction, we extended our AD recordings to include all pairwise combinations of head and scene velocities in the same set used above (Fig. 1k, Extended Data Fig. 2c). These extended recordings revealed the way in which the scene-velocity profile modulated the decoded head direction for a fixed head-velocity profile, over a range of head-velocity profiles (Fig. 1l).

We can distinguish two qualitatively different cases in the way visual and vestibular stimulations were combined. In the first case, the scene was rotated in the same direction as the head, and therefore the head and scene velocities had the same sign (Fig. 1l, right side). In this case, for each head speed the decoded head-direction shift was systematically reduced as compared to the case of head-only rotation (purple data points, center; 180 deg/s head speed and 45 deg/s scene velocity: p=2.5×10^-4^, in all other cases: p<10^-6^, one-sided paired t-test). As long as the scene speed was lower than the head speed, increasing the scene speed resulted in progressively smaller decoded head-direction shift. An intriguing situation was the one in which the scene speed exceeded the head speed (and visual motion indicated rotation in the direction opposite to the actual head rotation): we found that the decoded head-direction shift was sign-inverted with respect to the true angular head shift for a range of stimulation parameters (Extended Data Fig. 2d; p<0.01 for all such conditions, except for 135 deg/s head speed and 180 deg/s scene velocity, one-sided t-test for zero mean vs. negative mean). In other words, visual motion cues overrode the drive coming from vestibular motion cues. In the second case, the scene was rotated in the opposite direction to the head, and therefore the head and scene velocities had opposite signs (Fig. 1l, left side). Surprisingly, in this case a larger scene speed induced a significantly *smaller* shift in decoded head direction (p<0.05 for 11 of 16 pairwise comparisons; one-sided paired t-test). This is surprising because, for a simple, and in particular linear, combination of the motion cues in driving the decoded head-direction shift, one expects the opposite result.

### The slow phase of reflexive eye movement tightly correlates with the decoded head direction

In our experimental setup, we controlled the velocity of the scene relative to that of the head. However, visual motion signals to the brain arise from visual motion on the retina (“retinal slip”), which depends on scene motion, head motion, and eye movement relative to the head. To measure eye movement, we tracked the pupil of the right eye (7/10 mice), using an infrared camera, while recording neuronal population activity from AD (Fig. 2a). We expected to find eye movements resulting from the optokinetic reflex (OKR) and the vestibulo-ocular reflex (VOR)^30,38,39^. OKR and VOR evoke slow eye movements which stabilize the image on the retina, as well as corrective quick saccades which reset the position of the eye. We isolated the slow eye movements by removing saccades from the time trace of the eye velocity relative to the head. As a result, we obtained smooth curves of the angular shift of the eye for each experimental condition (Fig. 2b-d).

**Fig.2.**
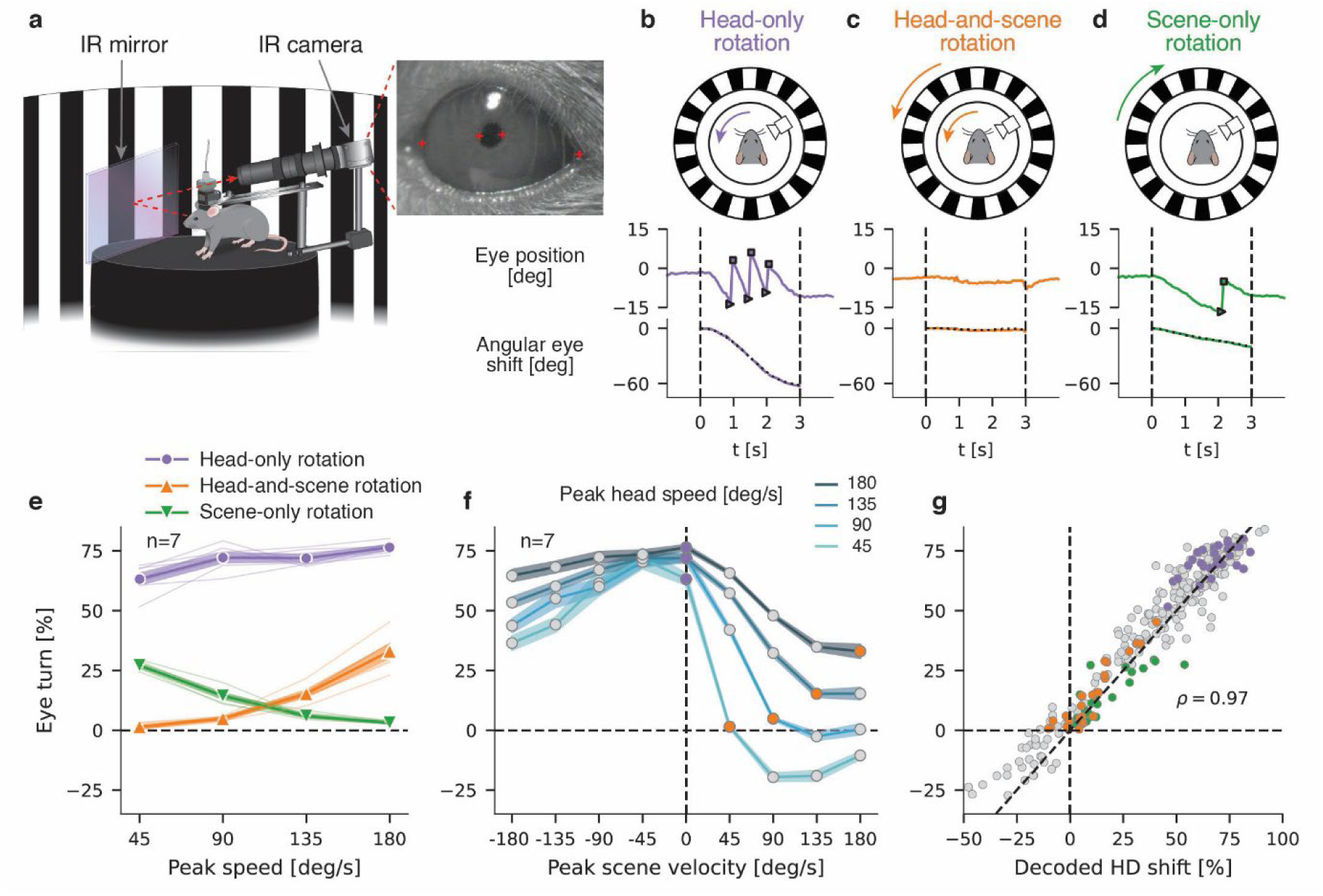
Tight correlation between slow eye movement and decoded head direction. **a**, Schematic of the experimental configuration for simultaneous recording of AD neuronal activity and measurement of eye movements in head-fixed mice during visual-vestibular stimulation. **b**, Top: Illustration of experimental design in which the eye position is tracked while the head rotates, and the scene remains stationary. Middle: Eye position over time for an example trial with a peak stimulus speed of 45 deg/s. Bottom: Angular shift of the eye derived from the slow eye movements, ignoring saccades, (dashed black line) and unwrapped eye position (solid colored line). **c** and **d**, Same as **b** but for experimental conditions in which both head and scene rotate in synchrony (**c**) or only the scene rotates (**d**). **e**, Eye turn—in percent normalized to the angular shift of either head (purple and orange) or scene (green)—as a function of peak stimulus speed in the three experimental conditions illustrated in **b**-**d** (n=7 mice). Thick lines and markers indicate the mean across animals (n=7), shaded areas indicate the standard error of the mean, and thin lines indicate individual mice. Since eye turns were highly similar for CW and CCW stimulus rotations with the same peak speed, we aggregated data from the two rotation directions the same way as done for the decoded head-direction shifts. **f**, Eye turn as a function of the peak scene velocity for each non-zero value of the peak head speed (n=7 mice). Purple and orange markers correspond to the same data as in **e**. **g**, Scatter plot of the eye turn vs. the decoded head-direction shift. Points correspond to average values per animal (n=7 mice). Markers are color-coded as in **e** and **f**. The dashed diagonal corresponds to the identity line.

We quantified the total angular shift of the eye during the stimulation. For convenience we inverted the sign of this shift to match that of the decoded head-direction shift. Henceforth, we refer to the resulting quantity as “eye turn.” Remarkably, in response to head-only, head-and-scene, and scene-only rotations, the eye turn normalized to the angular shift of head or scene was nearly identical to the corresponding decoded head-direction shift (compare Fig. 2e and Fig. 1j). This similarity held for all pairwise combinations of head and scene velocities (compare Fig. 2f and Fig. 1l, see also Extended Data Fig. 2e). As a result, the eye turn vs. the decoded head-direction shift datapoints lay close to the identity line and had a Pearson correlation of 0.97 (Fig. 2g; p=3.6×10^-170^, Pearson correlation test).

To understand the relation between eye movement and decoded head direction more precisely, we examined the time course of the angular shift of the eye and of the decoded head direction during stimulation (Extended Data Fig. 3). Interestingly, while the two quantities closely tracked each other in most experimental conditions, we also found discrepancies for a range of stimulation parameters, notably in situations in which the scene rotated in the same direction as and as fast or faster than the head (Extended Data Fig. 3). Such situations are not encountered in natural behavior. These systematic discrepancies were useful to refine a model of the neuronal computations giving rise both to an internal estimate of the head direction and to eye movement (see below).

### A model of visual-vestibular interaction explains the computation of head direction and reflexive eye movement

We aimed to develop the most parsimonious mathematical model which would both be consistent with what is known about visual and vestibular processing in the brain and explain our data quantitatively. We assumed that the decoded head direction was obtained by integrating a velocity signal^20,21^. The model’s task was to convert visual and vestibular motion cues to two velocity signals, one leading to the decoded head direction by integration, and the other corresponding to the angular velocity of the eye.

In the model (Fig. 3a), an estimate of the head velocity is obtained from a non-linear combination of a “visual signal” and a “vestibular signal.” The vestibular signal is obtained from a model of sensory transduction in the horizontal semicircular canals located in the inner ears^40^. The visual signal is a non-monotonic function of the retinal slip that mimics the sensitivity of visual neurons to motion, with activity suppressed both at low and high speeds. This form agrees with the response of direction-selective retinal ganglion cells as a function of speed^41^, and is in line with classic models of the OKR^42,43^. In turn, retinal slip is computed as a linear combination of scene motion, head motion, and eye movement with respect to the head. Scene motion and head motion are the inputs to the model. A crucial proposal in the model is that eye movement is governed in large part by the model’s head velocity estimate. This is a natural assumption: in the case of a globally stationary visual world, reflexive eye movements stabilize the image on the retina during head motion. The assumption is also motivated by the strong correlation between the eye turn and decoded head-direction shift in the data. Therefore, the model connects head-direction coding to reflexive eye movement and elucidates their coupling. The observation of few discrepancies between the decoded head direction and the angular eye shift for specific types of head- and scene-motion stimulation allowed us to refine the coupling between the two systems. We found that a small, direct contribution of the vestibular signal to the eye motor command was needed to capture eye movement quantitatively, particularly for the conditions in which we observed discrepancies between the decoded head direction and angular eye shift. Finally, the model takes into account the delay of eye movements with respect to motor commands due to the biomechanical properties of the eye^44,45^.

**Fig.3.**
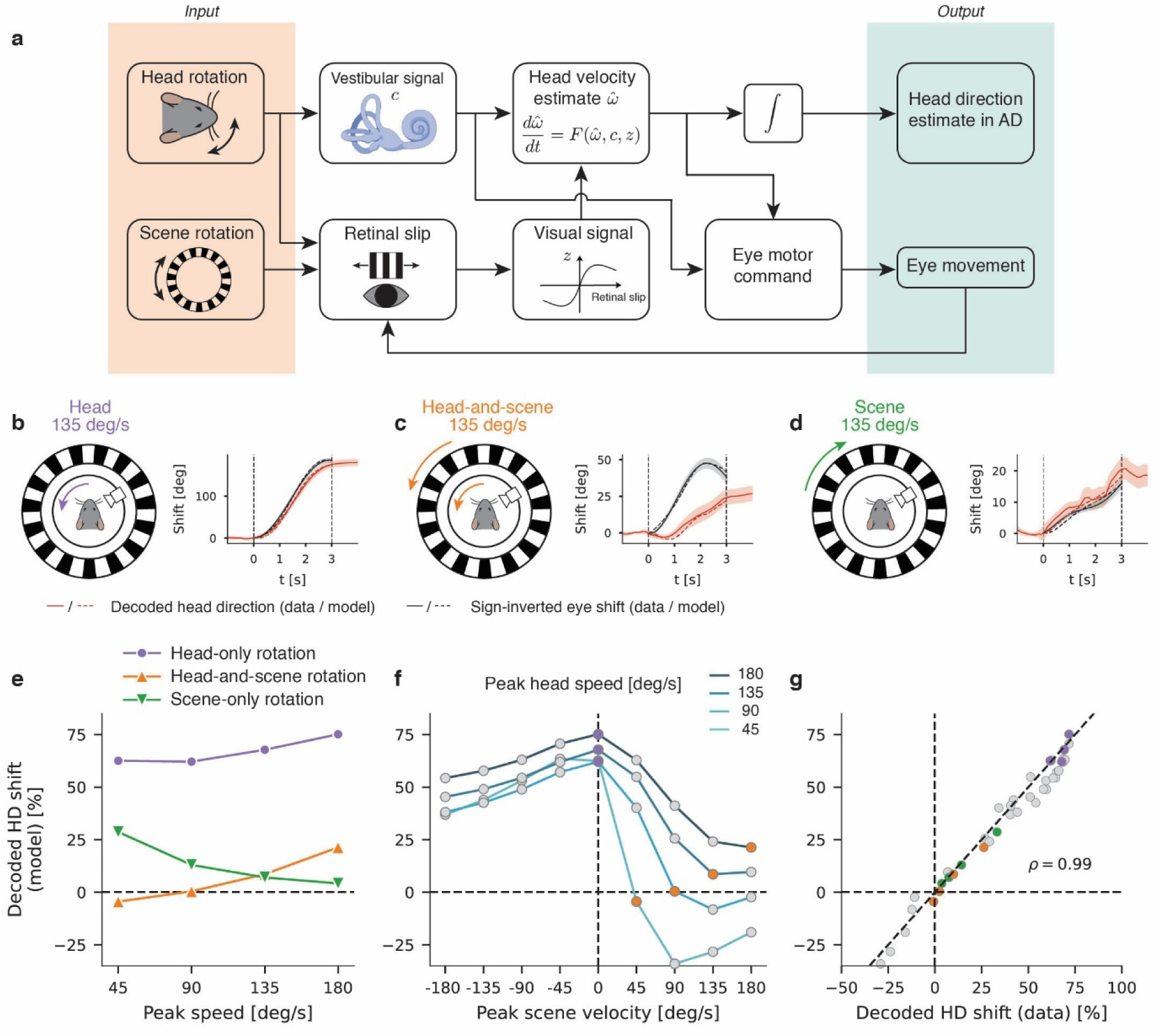
Model of the joint computation of head direction and slow eye movement. **a**, The head-direction estimate results from integration of an estimate 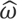 of angular head velocity in the head-direction system. The time derivative of 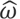 is a function of 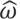 itself, a vestibular signal *c* and a visual signal *z*. The vestibular signal *c* is generated in the semicircular canals in response to rotations of the head. The visual signal *z* is derived from the retinal slip velocity *r* through a nonlinear transformation. Retinal slip velocity *r* is the sum of the scene velocity (which is usually zero in natural conditions) and the additive inverse of eye-in-head velocity and head-in-world velocity. Finally, both the estimate 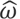 and the vestibular signal *c* are used to generate eye movement. **b**, Left: Illustration of the experimental design in which the head rotates with a peak speed of 135 deg/s while the scene remains stationary. Right: Time series of the decoded head-direction shift (red lines), the sign-inverted angular shift of the eye (black lines) for data (solid lines) and model (dashed lines). Data traces show the average across animals (n=7 mice) after alignment of CW and CCW data. The shaded area indicates the standard error of the mean. **c** and **d**, Same as **b** but for experimental conditions in which both head and scene rotate in synchrony (**c**) or only the scene rotates (**d**). **e** and **f**, Normalized decoded head-direction shift predicted by the model for the different experimental conditions. **g**, Scatter plot of the predicted vs. the measured decoded head-direction shift. Markers are color-coded as in **e** and **f**.

The model reproduced the time courses of both the decoded head direction and the angular eye shift with quantitative accuracy (Fig. 3b-d and Extended Data Fig. 3). In particular, it captured the dependence of both decoded head-direction shift and eye turn on the peak stimulus speed for head-only, head-and-scene, and scene-only rotations (Fig. 3e and Extended Data Fig. 4a). More broadly, the model reproduced the behavior of the decoded head-direction shift and the eye turn for all pairwise combinations of head and scene velocities (Fig. 3f and Extended Data Fig. 4b, compare with Fig. 1l and Fig. 2f). A plot of the model prediction of the decoded head-direction shift vs. the measured decoded head-direction shift, for all experimental conditions, further confirmed the ability of the model to capture the data quantitatively, with a Pearson correlation of 0.99 (Fig. 3g). Furthermore, sensitivity analyses indicated that removing terms in the model or changing parameter values degraded its performance (Extended Data Fig. 4c-f), suggesting that the model does not contain redundant components.

### Horizontal direction-selective retinal ganglion cells are the source of visual motion signals for the head-direction system

A central element of the model is that the OKR and the head-direction system both rely on the same visual signal. The model therefore predicts that selective perturbation of the visual circuitry underlying the OKR disrupts not only reflexive eye movement, but also the instant-to-instant updating of head direction. Specifically, if a perturbation of the visual system abolishes the OKR, then the predicted decoded head-direction shift is obtained by setting the visual signal in the model to zero (Fig. 4a). Under this condition, the model predicts impaired estimates of head-direction shift, as compared to predictions of the full model (in which the visual signal is not set to zero): the predicted decoded head-direction shift is vanishing at low head speed, and even though it increases with speed, it never reaches the magnitude obtained in the case of the full model (Fig. 4b). Furthermore, when the visual signal is set to zero in the model, the decoded head-direction shifts predicted for head-only rotation and head-and-scene rotation are the same (Fig. 4b). More generally, the model predicts that the decoded head-direction shift is not modulated by scene velocity at any given head speed (Fig. 4c).

**Fig.4.**
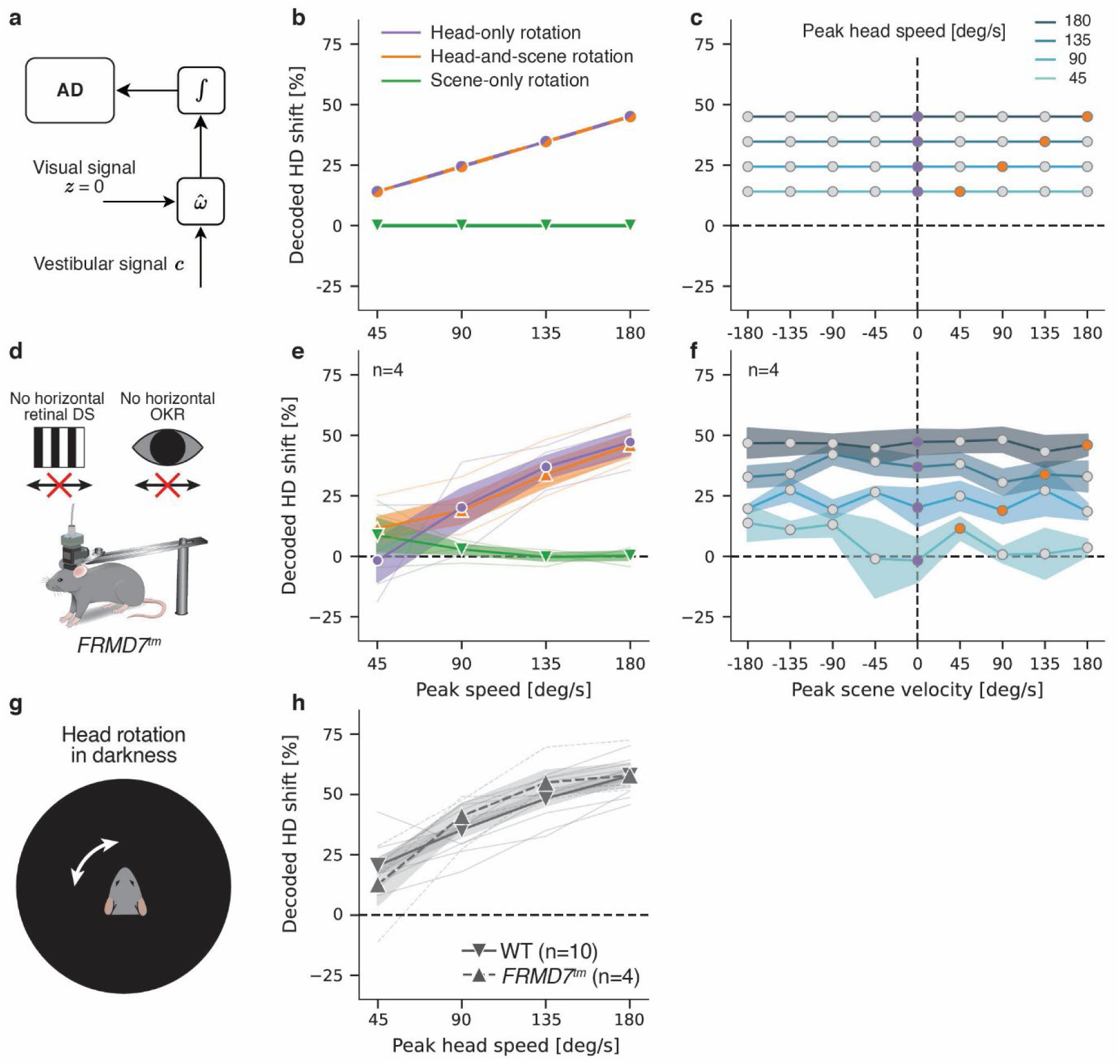
Horizontal direction-selective retinal ganglion cells as the source of visual motion signals to the head-direction system. **a**, When setting the visual signal to zero in the model, the shift of the AD head-direction estimate depends on the vestibular signal only. **b** and **c**, Model prediction for the decoded head-direction shift in the different experimental conditions with the visual signal set to zero. **d**, Calcium imaging in head-fixed *FRMD7^tm^* mice. *FRMD7^tm^* mice lack both horizontal direction selectivity (DS) in the retina and the horizontal optokinetic reflex (OKR). **e**, Decoded head-direction shift measured in *FRMD7^tm^* mice as a function of the peak stimulus speed in the following three experimental conditions: the head rotates while the scene remains stationary (purple), head and scene rotate in synchrony (orange), the scene rotates while the head remains stationary (green). Thick lines and markers indicate the mean across animals (n=4), shaded areas indicate the standard error of the mean, and thin lines indicate individual mice. The decoded head-direction shift, in percent, is normalized to the angular shift of either the head (purple and orange) or the scene (green). **f**, Decoded head-direction shift as a function of the peak scene velocity for each non-zero value of the peak head speed. The decoded head-direction shift is normalized to the actual head-direction shift. Shaded areas indicate the standard error of the mean (computed across n=4 mice). **e** and **f** aggregates data from CW and CCW rotations the same way as in Fig. 1. **g**, Schematic of head rotations in darkness. **h**, Decoded head-direction shift (in percent, normalized to the actual head-direction shift) as a function of peak head speed for wild-type (n=10) and *FRMD7^tm^* (n=4) mice. Thick lines and markers indicate the mean across animals, shaded areas indicate the standard error of the mean, and thin lines indicate individual mice.

The horizontal OKR is triggered by activity of retinal ganglion cells in response to horizontal visual motion^46^. Mice with a targeted mutation in the *Frmd7* gene (*FRMD7^tm^* mice) lack horizontal (but not vertical) direction-selectivity in the retina, and consequently lack the horizontal (but not the vertical) OKR^33^. Therefore, we hypothesized that in *FRMD7^tm^* mice the decoded head-direction shift would match the prediction of the “truncated model” in which the visual signal is set to zero.

We thus repeated the same head-fixed experiment with visual-vestibular stimulations as above, but with *FRMD7^tm^* mice (Fig. 4d). The decoded head-direction shift in *FRMD7^tm^* mice closely matched the model’s predictions (Fig. 4e,f). The decoded head-direction shift in head-only rotation was considerably lower at all speeds in *FRMD7^tm^*mice than in wild-type mice (Fig. 4e, compare with Fig. 1j; p<10^-4^ for all speeds, one-sided two-sample t-test). Furthermore, the decoded head-direction shift in *FRMD7^tm^* mice did not significantly differ between head-only and head-and-scene rotations at any speed (two-sided paired t-test). In addition, and in contrast to wild-type mice, in *FRMD7^tm^*mice, the decoded head direction did not significantly change during scene-only rotation at any of the speeds tested (Fig. 4e, compare with Fig. 1j; one-sided t-test for zero mean vs. positive mean). More broadly, we did not observe any significant modulation of the decoded head-direction shift by scene velocity for any of the head speeds we tested (Fig. 4f; two-sided paired t-test comparing conditions with non-zero scene speed against the head-only rotation condition). These results suggest that the same horizontal direction-selective retinal ganglion cells that drive the OKR are the source of visual signals used by the head-direction system, in the way predicted by a truncated model in which the visual signal is set to zero.

As an additional way to test the model’s prediction in the absence of visual motion signals, we performed recordings during head rotation in complete darkness (Fig. 4g). The corresponding decoded head-direction shift exhibited no significant difference between wild-type and *FRMD7^tm^*mice at all speeds tested (two-sided two-sample t-test), and closely matched model predictions (Fig. 4h and Extended Data Fig. 5a). The model slightly but systematically underestimated the decoded head-direction shift for all values of the head speed. This discrepancy can be modeled by assuming a light-induced modulation of a gain factor in the model, with a greater vestibular gain in darkness (Extended Data Fig. 5b). In summary, under two separate perturbations of the visual system, *FRMD7^tm^* mice and complete darkness, the model accurately predicted the decoded head-direction shift in the absence of visual motion signals, in head-fixed conditions.

Interestingly, when we recorded from head-direction cells in freely moving *FRMD7^tm^* mice in the presence of visual landmarks, we could decode head direction as well as in wild-type mice (Extended Data Fig. 6; see also Extended Data Fig. 1h). This suggests that either other visual signals or self-motion signals such as proprioceptive and motor signals, present during development, could substitute for absent visual motion signals from horizontal direction-selective retinal ganglion cells.

### Perturbation of visual motion cues alters navigation behavior

While head-fixed experiments allowed for the separate control of visual and vestibular motion cues, it remained an open question whether visual motion cues also affect the updating of the head-direction estimate in freely moving mice. We therefore recorded from the AD in freely moving wild-type and *FRMD7^tm^* mice, and we briefly rotated the scene at random times (Fig. 5a and Supplementary Video 1). As before, the scene consisted of vertical contrast bars displayed on the cylindrical LED screen, and rotations of the scene lasted for three seconds, with a peak speed of 45 deg/s. When comparing the decoded and the actual head-direction shifts in wild-type mice, we observed that CW scene rotations biased the decoded head-direction shift CCW, and vice versa for CCW scene rotations, as expected from the findings in head-fixed mice (Fig. 5b). To assess the robustness of this bias in freely moving wild-type mice, we computed the average difference between the decoded and the actual head-direction shifts (Δ) for each animal. We found a significant difference in Δ between the CW and the CCW scene rotations (Fig. 5c; p=3.8×10^-5^, one-sided paired t-test). By contrast, and as expected from the findings in head-fixed mice, scene rotations did not bias the decoded head-direction shift in *FRMD7^tm^* mice (Fig. 5d-f; one-sided paired t-test on Δ: non-significant). Thus, visual motion cues affected the decoded head direction in freely moving wild-type mice, and this depended on intact horizontal direction-selectivity in retinal ganglion cells.

**Fig.5.**
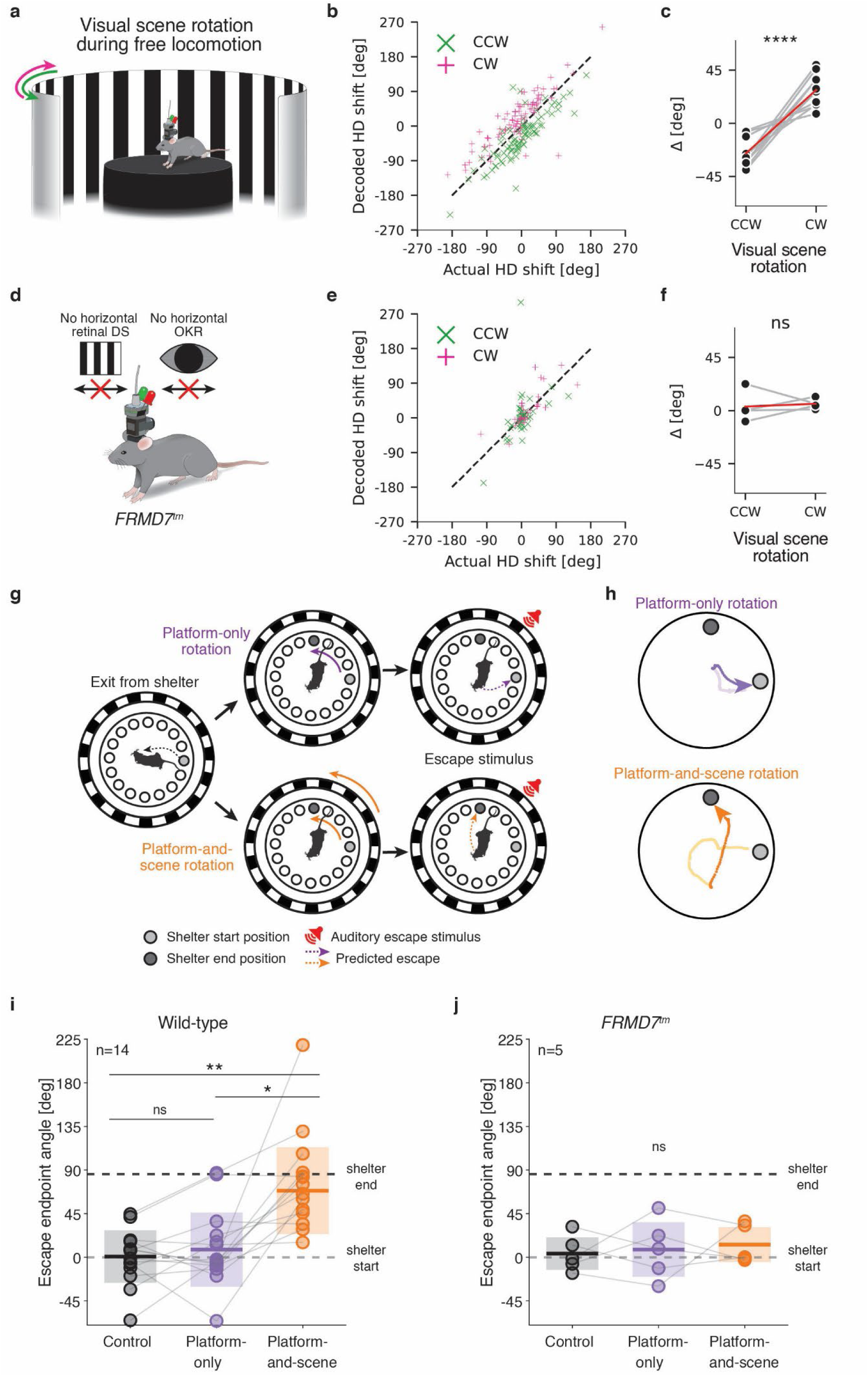
Alterations of mouse navigational behavior from perturbations of visual motion cues. **a**, Calcium imaging in the AD of freely moving mice. At random intervals the scene was rotated either CW or CCW for 3 seconds with 45 deg/s peak speed. **b**, Decoded head-direction shift (degrees) vs. actual head-direction shift (degrees) during every scene rotation trial for all animals (300 trials from n=10 mice). Pink markers correspond to CW scene rotations, green markers to CCW scene rotations. Positive values indicate CCW shifts of actual and decoded head direction, negative values indicate CW shifts of actual and decoded head direction. **c**, The difference between the angular shifts of decoded and actual head direction during scene perturbation trials, averaged across all CW and CCW trials of the recording for each mouse (black data points and grey lines) and averaged across mice (red line). t-test on related samples: **** = p<0.0001. **d**, Schematic of recording from *FRMD7^tm^*mice. **e**, Same as **b** but for *FRMD7^tm^* mice. **f**, Same as **c** but for *FRMD7^tm^* mice. t-test on related samples: p>0.05. **g**, Schematic of experiment. A trial is initiated by the mouse exiting the shelter and walking to the center of the platform. Next, one of two perturbations took place: either only the platform was rotated for 3 seconds with a peak speed of 45 deg/s (top), or both the platform and the scene were rotated in synchrony for 3 seconds with a peak speed of 45 deg/s (bottom). Immediately after the perturbation, an auditory escape stimulus was presented. In control trials (not shown in the schematic), the escape stimulus was presented after the mouse exited the shelter and walked to the center of the platform, with no platform or scene perturbation. **h**, Example trajectories during the two different perturbation trials for one mouse. Pale colors indicate the position of the mouse (center of mass) at each time point before the escape stimulus presentation, darker colors indicate the position at each time point after the escape stimulus presentation. Note that for analysis and visualization, the coordinate system was rotated such that the shelter start position is at zero degrees. Data from CW and CCW perturbations were aligned. **i**, The angle of the endpoint of the escape response (in degrees, relative to the shelter start position where the shelter start position is 0 degrees) across conditions. Connected circles represent individual mice, solid horizontal lines represent the circular mean across mice for the given condition. The shaded boxes show the circular standard deviation. Wilcoxon signed-rank tests on the sine of pairwise differences, with Bonferroni-Holm correction for multiple comparisons: ns = p>0.05, * = p<0.05, ** = p<0.01. **j**, Same as **i** but for *FRMD7^tm^*mice. Wilcoxon signed-rank tests on the sine of pairwise differences, with Bonferroni-Holm correction for multiple comparisons: p>0.05 for all comparisons.

Since the head-direction system is known to support spatial navigation^9,10^, we asked whether visual motion cues also subserve spatial navigation. To answer this question, we devised a behavioral paradigm which makes use of an innate defensive behavior in mice: running to a shelter when presented with an aversive auditory stimulus^47,48^ (Fig. 5g). Mice (n=14) were freely moving on a platform containing 16 equally sized, equidistant holes around the edge of the platform. One of these holes was an entrance to a shelter hidden below the platform, while all the other holes were sealed with transparent plexiglass, and served as decoys. We opted for this design because it was shown before that mice rapidly learn a shelter’s location and use path integration, as opposed to landmark-based navigation, to escape a threat by running to the shelter^48^. We examined whether perturbing visual motion cues before the presentation of an aversive sound alters the escape directions taken by freely moving mice.

There were three different types of trial in our experiments (with one trial per behavior session): control trial, “platform-only rotation” trial, and “platform-and-scene rotation” trial. The last two conditions were analogues to the head-only rotation and head-and-scene rotation conditions in head-fixed mice. During a baseline period, before any trial, we allowed the mouse to spontaneously find the shelter and learn entering and exiting the shelter. Trials were triggered when the mouse walked to the center of the platform after spontaneously exiting the shelter. In a control trial, once the mouse reached the center of the platform, an auditory stimulus was played and elicited an escape response. In a platform-only rotation trial, once the mouse reached the center of the platform, it was rotated either CW or CCW, with a peak speed of 45 deg/s; then, immediately after the platform rotation, the auditory stimulus was played. The platform-and-scene rotation trial only differed in that the scene was rotated together with the platform.

We expected that mice correctly account for the rotation when deciding on their escape trajectory in platform-only rotation trials, since, in head-fixed experiments, head-only rotation significantly shifted the decoded head direction (Fig. 1j, purple curve). Conversely, we expected that mice would fail to account for the rotation in platform-and-scene rotation trials, and therefore display a biased escape trajectory. This is because, in head-fixed experiments, head-and-scene rotation at 45 deg/s speed did not shift the decoded head direction (Fig. 1j, orange curve). Note that, since the shelter was physically attached to the platform, an escape trajectory which *does* account for the rotation of the platform would end up at the pre-rotation location of the shelter (“shelter start position”), while an escape trajectory that does *not* account for the rotation of the platform would end up at the post-rotation location of the shelter (“shelter end position”). Therefore, we predicted that mice would escape to (approximately) the shelter start position in the platform-only rotation trials, and to the shelter end position in the platform-and-scene rotation trials.

To quantify escape trajectories, we defined an “escape angle” as the angle between the vector connecting the platform center to the position of the head of the mouse at the endpoint of the escape, and the vector connecting the platform center to the center of the shelter start position (Extended Data Fig. 7). To combine CW and CCW trials, we inverted the sign of escape angles in CW trials. An escape angle of zero degrees corresponds to an escape to precisely the shelter start position, while an escape angle of 85.9 degrees (the angle by which the platform and shelter were rotated) corresponds to an escape precisely to the shelter end position. Control trials served to ascertain that mice correctly orient and, as expected, the circular mean of the escape angle was close to zero (0.7±27.1 deg, circular mean ± circular standard deviation) (Supplementary Video 2). In platform-only rotation trials, the circular mean of the escape angle was 7.9±38.3 deg (Fig. 5h,i; Supplementary Video 3), close to the shelter start position (0 deg) and did not significantly differ from control trials (two-sided Wilcoxon signed-rank tests on the sine of the difference between the escape angles), suggesting that mice accounted for the rotation of the platform when both visual and vestibular motion cues indicated rotation. By contrast, in platform-and-scene rotation trials, the circular mean of the escape angle was 68.7±44.8 deg (Fig. 5h,i; Supplementary Video 4), close to the angle of the shelter end position (85.9 deg), and significantly different from platform-only (p=0.01) and control (p=9.2×10^-3^) trials. This suggests that mice did not account for the rotation of the platform when only vestibular motion cues indicated rotation. These results demonstrate that visual motion cues profoundly impact spatial navigation in freely moving mice, in agreement with both the findings in head-fixed mice and the mathematical model of the visual-vestibular process updating the head-direction estimate.

To test whether the ability to use visual motion cues for spatial navigation depends on horizontal direction-selective retinal ganglion cells we repeated the spatial navigation task with *FRMD7^tm^*mice. Consistent with the observation that freely moving *FRMD7^tm^*mice have an intact head-direction representation (Extended Data Fig. 6), but are not impacted by visual motion cues (Fig. 5d-f), *FRMD7^tm^* mice escaped toward the shelter start position in all three conditions (Fig. 5j; two-sided Wilcoxon signed-rank tests on the sine of the difference between the escape angles: no significant differences). This indicates that *FRMD7^tm^* mice account for the rotation of the platform without the use of visual motion cues. Taken together, these results demonstrate that horizontal direction selective ganglion cells are the source of visual motion signals used for goal-directed spatial navigation in freely moving mice.

### Statistically optimal estimation of head direction

During free locomotion, the brain likely uses not only visual and vestibular motion signals but also motor signals to update its head-direction estimate. Previously in the paper, we proposed a mechanistic model showing how visual and vestibular motion signals interact under head-fixation—when motor signals are absent—to form a multisensory velocity estimate used to update the head-direction estimate. Here we asked what the predictions of a statistically optimal estimation for such a situation would be, that aims to keep track of angular head velocity from a continuous stream of noisy self-motion signals. As in our mechanistic model, we assumed that the visual signal results from a nonlinear transformation of the retinal slip, that eye motion is driven by the head-velocity estimate and that the vestibular signal arises from sensory transduction in the inner ear. In addition, we introduced a motor signal that, under natural conditions, provides an unbiased estimate of angular head velocity. We further assumed that the brain generally expects head motion to be accompanied by such a motor signal, also in the unnatural head-fixed condition. All sensory signals were assumed to be noisy (Extended Data Fig. 8). Bayes filtering offers a statistically optimal approach to estimating unknown quantities from such noisy signals. In our case, the Bayes filter computes a probability distribution over angular head velocity, given all past and the current multisensory signals (Fig. 6a). We examined the filter’s predictions for the head and scene rotation stimuli applied in our experiments, corresponding to the unnatural situation in which motor signals are absent, i.e., set to zero. Strikingly, the predictions for the decoded head-direction shift closely matched our experimental observations (Fig. 6b,c, Extended Data Fig. 9). The underestimation of the head-direction shift during head-only rotations is explained by the lack of motor input. This suggests that our findings in head-fixed mice reflect a multisensory velocity estimation process that is optimized for the natural condition, where visual and vestibular motion cues are congruent, and additional motor signals are available.

**Fig.6.**
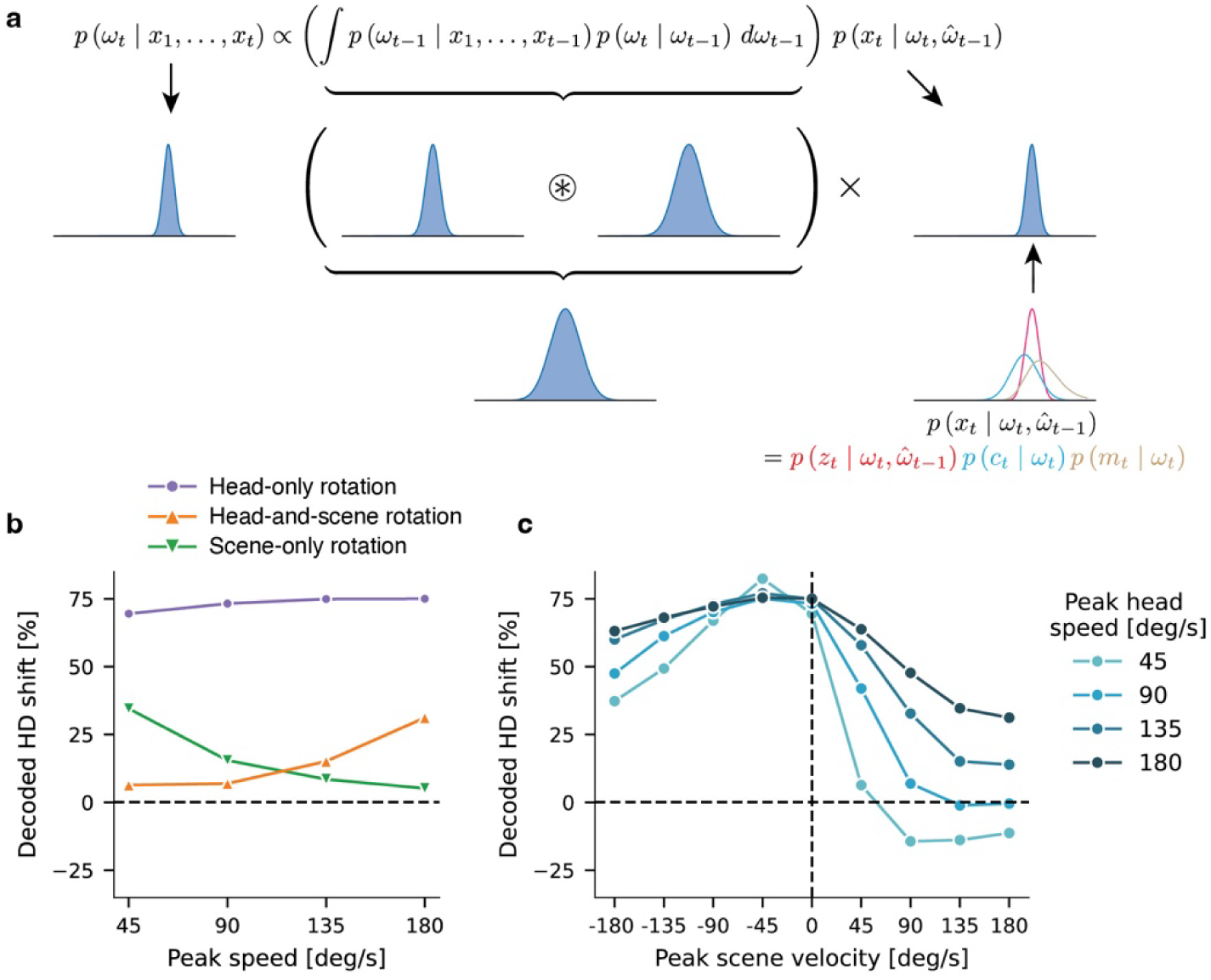
Bayesian model of multisensory estimation of angular head velocity. **a**, A Bayes filter computes the posterior probability distribution *p*(ω_*t*_ | *x*_1_, …, *x*_*t*_) over the angular head velocity ω*ₜ* given sensory inputs *x*_1_, …, *x*_*t*_. The computation is done in a sequential manner, whereby the posterior distribution at the previous time step is first convolved with the transition probability *p*(ω_*t*_ | ω_*t*−1_). The resulting (predictive) probabilities are then multiplied with the likelihood 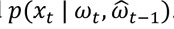. The latter factorizes into the likelihoods of the visual signal *z*_*t*_, the vestibular signal *c*_*t*_ and the motor signal *m*_*t*_; the visual signal depends on the eye velocity, set equal to the posterior mean 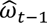 at the previous time step. **b** and **c**, Normalized decoded head-direction shift obtained from the Bayesian model for the different experimental conditions under head fixation in brightness. The decoded head-direction shift corresponds to the temporal integral over the posterior mean head velocity. In the simulations, the visual signal and the vestibular signal were set to their mean values, given the current head velocity and the posterior mean from the previous time step; the motor signal was set to zero.

## Discussion

Our results demonstrate that both the brain’s estimate of head direction and goal-directed navigation critically depend on visual motion cues. Specifically, visual motion cues contribute to the instant-to-instant updating of the head-direction estimate in AD. While vestibular cues dominate at high speeds, visual cues dominate at low speeds; importantly, however, neither cue alone results in an adequate head-direction estimate, at any speed we tested. A very slow rotation of a visual point array (4.5 deg/s) presented over a prolonged time was previously found to influence head-direction cell activity^28^. Our work disentangles the contributions of visual and vestibular motion cues over a range of behaviorally relevant speeds, and quantifies the impact of transient visual motion cues on the head-direction system. We have furthermore shown that visual motion cues are signaled via direction-selective retinal ganglion cells.

We found that, across a wide range of passive visual-vestibular stimulation conditions, the head-direction shift decoded from AD activity is almost identical to the eye turn. Both the decoded head-direction shift and the eye turn most closely followed the actual head-direction shift in the “natural” condition where the visual scene is stable. This suggests that reflexive slow eye movements controlled by the oculomotor system are adapted to stabilize the retinal image of a naturally static world. In accordance, a large fraction of eye movements observed in freely moving mice stabilize gaze during rotations of the head around its yaw axis^49,50^. In this study, we focused on the slow phase of reflexive eye movement in head-fixed animals. Future work should investigate the nature of the coupling between the head-direction system and the oculomotor system in freely moving animals, when motor efferent signals and different types of eye movements are present. Interestingly, the direction and amplitude of saccades also correlate with changes of the head-direction estimate in the AD of freely moving mice^51^.

The tight coupling of head-direction estimate and reflexive eye movement, as well as the shared reliance on visual inputs from direction-selective retinal ganglion cells, suggest that a cognitive function, namely head-direction estimation, and reflexes, namely VOR and OKR, share neuronal circuitry and computation. As VOR and OKR are highly conserved in vertebrates^52^, and head-direction networks across species share key properties^53^, we expect these findings in mice to extend to other mammalian species.

To understand the non-trivial combination of visual and vestibular signals in the brain’s estimate of head direction, and to capture its tight coupling with eye movement and their discrepancy in some experimental conditions, we proposed and validated a computational model that predicts both the head-direction estimate and the slow phase of reflexive eye movement from visual and vestibular cues. The model explained our experimental observations not only in wild-type mice, but also in *FRMD7^tm^* mice which selectively lack horizontal direction-selectivity in retinal ganglion cells and consequently lack the horizontal OKR. In *FRMD7^tm^* mice, visual motion cues failed to update the head direction representation both during head-fixation and free movement. Furthermore, perturbation of visual motion cues impacted goal-directed spatial navigation only in wild-type mice but not in *FRMD7^tm^* mice. These results demonstrate that the visual motion signals conveyed by horizontal direction-selective ganglion cells are used to drive “low-level” reflexive behavior (OKR) as well as “high-level” cognitive behavior (spatial navigation).

We examined the combination of visual and vestibular signals under head-fixation, and found that the decoded head-direction shift systematically underestimated the actual head-direction shift in response to head-only rotations under these conditions. We interpret this discrepancy as evidence that, in addition to the crucial roles of visual and vestibular motion signals, the brain also integrates motor signals, available during free locomotion, into its velocity estimation process. Supporting this interpretation, a probabilistic estimation framework that incorporates the motor modality accounts well for the measurements obtained under head-fixation. The core assumptions of the probabilistic approach are aligned with our mechanistic model, particularly the tight coupling between eye movements and the head-direction estimate. These results suggest that our findings in head-fixed mice reflect a multisensory velocity estimation process that is optimized for the natural condition, where visual and vestibular signals are congruent, and motor signals are also available. Taken together, our data is well-described by a mechanistic model, which elucidates how visual and vestibular signals are integrated to drive both the updating of the head-direction representation and reflexive eye movement, as well as a probabilistic model which indicates that visual and vestibular signals are combined in a statistically optimal manner.

## Acknowledgements

We thank the Complex Viruses Platform for AAV production; present and past members of the Roska lab for discussions and comments; G. Kosche and A. Fratzl for discussions and technical assistance; F. Franke, A. Fratzl, G. Kosche, and T.M. Rodrigues for comments on the manuscript; SciDraw (doi.org/10.5281/zenodo.3925917) and V. Juvin (SciArtWork) for graphics. Part of the calculations were performed at sciCORE (http://scicore.unibas.ch/) scientific computing center at University of Basel. This study was funded by Sedinum Foundation fellowships to T.D. and L.E., Swiss National Science Foundation Synergia grant (CRSII3_141801), European Research Council advanced grant (HURET N°883781), Carigest grant, Louis-Jeantet Foundation award, Körber Foundation award, Swiss National Science Foundation grant (31003A_182523), and the NCCR ‘Molecular Systems Engineering’ to B.R., and the European Research Council (ERC) under the European Union’s Horizon 2020 research and innovation program grant agreement No. 866376 to R.A.S.

## Author contributions

Conceptualization, Experiment design, Writing: T.D., L.E., B.R., R.A.S. Investigation, Data Curation, Formal analysis, Software, Visualization: T.D., L.E. Model: L.E., R.A.S. Supervision, Funding acquisition: B.R., R.A.S.

Equally-contributing authors ordered alphabetically.

## Correspondence and requests for materials

Correspondence should be addressed to Botond Roska or Rava Azeredo da Silveira.

## Data availability

The data reported in this paper is available from the corresponding authors upon request.

## Code availability

Custom code used for analysis is available at https://github.com/l-ew/vis-vest-hd-computation.

## Methods

### Animals

Female mice were used for experiments. Bl6 mice (“wild-type”, strain: C57BL/6J) were purchased from Charles River laboratories, *FRMD7^tm^* mice (FRMD7tm1a(KOMP)Wtsi) were obtained from the Knockout Mouse Project (KOMP) Repository and backcrossed with Bl6 mice^33^. Mice were aged 8-12 weeks at the time of surgery and aged 11-25 weeks during the recordings. Mice in the behavioral experiment were aged 8-10 weeks old at the time of the experiment. Mice were group-housed and maintained on a normal 12-hour light/dark cycle with ad libitum access to food and water. Animal experiments were performed in accordance with standard ethical guidelines (European Communities Guidelines on the Care and Use of Laboratory Animals, 86/609/EEC) and were approved by the Veterinary Department of the Canton of Basel-Stadt.

### Surgeries

Mice were anesthetized with a Fentanyl-Medetomidine-Midazolam (FMM) mix: Fentanyl (Janssen, 0.05 mg/kg), Medetomidine (Virbac AG, 0.5 mg/kg), Midazolam (Sintetica, 5 mg/kg). The scalp was shaved with a clipper (Aesculap, Isis) and mice were placed on a 38 °C heat pad and fixed in a stereotaxic frame (Model 1900, Kopf). Eye gel (Coliquifilm, Allergan) was applied to prevent corneal dehydration. The scalp was disinfected and local anesthetic (Bupivacaine (0.25%) + Lidocaine (0.1-0.2%)) was injected subcutaneously. The scalp was then removed, and the skull cleared of connective tissue. To target the anterodorsal thalamus in the left hemisphere, a burr hole was made using a 0.8 mm diameter carbide drill bit (McMaster-Carr) in a stereotaxic drill (Model 1911, Kopf), 0.78 mm lateral to and 0.9 mm posterior to bregma. A sterile 23G syringe was lowered 2.1 mm into the brain to create a tract prior to AAV injection and GRIN lens implantation. 100-150 nl of AAV2/1-EF1a-GcaMP6s was injected between -2.9 mm and -2.7 mm below the brain surface using a calibrated glass pipette (Drummond Scientific Company) connected to a pressure ejection system (PDES-02DX, NPI). A GRIN lens (0.6 mm diameter, 7.3 mm length) with the microscope baseplate already attached (ProView Integrated Lens, Inscopix) was lowered to -2.78 mm below the brain surface using a custom holder and fixed to the skull using dental cement (Superbond C&B). A custom titanium head-bar was also attached with dental cement. At the end of the surgery an antagonist mixture, consisting of Flumazenil (0.5 mg/kg) and Atipamezol (2.5 mg/kg), was injected to counteract FMM anesthesia, and Buprenorphine (0.05-0.1 mg/kg) was injected for analgesia. Mice were monitored daily following surgery and allowed to recover for at least 4 weeks before any recordings.

### Experiment arena

We built a custom experiment arena which allowed for precisely controlled visual and vestibular stimuli. The arena consisted of a 360-degree cylindrical LED display with 96.5 cm height, 100 cm inner diameter, and 2 mm pixel pitch (Shenzen Bescanled), which was controlled by a Windows PC via an LED video processor (MSD300, Novastar). Visual stimuli were generated and presented using PsychoPy^54^. In the center of the cylinder, we built a central enclosure (48 cm height, 32 cm diameter) which housed a brushless DC motor (D5065 270KV, ODrive) connected to a 50:1 planetary reducer (PLX060, Yun Duan, AliExpress), which in turn was connected to a circular breadboard (20 cm diameter, Base Lab Tools) via a custom adapter plate. Different platforms were attached to the breadboard for different experiments. The DC motor was connected to an encoder (CUI AMT10E2-V) and motor control board (ODrive v3.6), which allowed custom control of the motor. All the cables from the motor ran internally from the central enclosure through the wooden floor of the arena, which was raised relative to the floor of the room, meaning that no cables were visible on the floor of the arena. The top of the arena was covered with a sheet of blackout fabric (BK5, Thor Labs), with a small cut-out in the center allowing for cables from the head-mounted microscope and eye tracking camera to pass through, as well as providing space for a camera to record the mouse from above.

### Data acquisition

In vivo calcium imaging was performed with a head-mounted miniature microscope (nVista 3.0, Inscopix). The microscope cable was attached to a commutator (Inscopix) to reduce torsional forces on the cable, which in turn was connected to the microscope data acquisition box (DAQ; Inscopix). The DAQ was connected to the local area network (LAN) with an ethernet cable, and controlled by the Inscopix Data Acquisition Software in a Google Chrome browser running on a MacBook Pro. For acquisition of calcium imaging videos, the excitation LED (475 nm) was set to 0.5 – 1.4 mW/mm^2^ and the gain adjusted depending on the mouse to maximize fluorescence signal quality. The electronic focus was set to 550 – 1000 in the Inscopix Data Acquisition Software (corresponding to 165 – 300 µm) depending on the mouse to maximize the number of cells in focus. Calcium imaging videos were acquired at a frame rate of 12 Hz, with each frame being timestamped by the DAQ. Top-view videos of the mouse during each recording session were acquired using a camera (a2A2590-60ucPRO, Basler; wide angle lens #89-524, Edmund Optics) mounted centrally above the platform at a frame rate of 30 Hz. In experiments in which the animal was moving freely as well as in head-fixed experiments in brightness and without eye-tracking, a red (700 nm, Kingbright) and a green (565 nm, Kingbright) LED were attached to the microscope to enable tracking of the head direction in the videos recorded with the top-view camera. In most of the experiments with head-fixation, to enable pupil tracking, we recorded videos of the right eye at a frame rate of 60 Hz using a camera (a2A 1920-160ucPRO, Basler; VZM™ 200i Zoom Imaging Lens, Edmund Optics) mounted behind and to the left of the mouse. To enable visibility of the pupil, the eye was illuminated from above using an 850 nm LED array (Thor Labs) with a longpass glass filter (cut-on wavelength: 830 nm, Schott RG830, Edmund Optics) attached in front of it. The image of the eye was reflected to the camera from a transparent hot-mirror (101.6×127 mm, 0 degree angle of incidence, Edmund Optics) placed in front of the eye. The frames of the top-view and eye tracking videos were timestamped by sending the exposure status of both cameras to the microscope DAQ via BNC cables. Top-view and eye tracking videos were subsequently aligned to the calcium imaging videos via linear interpolation. During experiments under head-fixation, timestamps of trial onsets and offsets were sent from a LabJack (LabJack U3-LV) to the microscope DAQ via a BNC cable. Prior to recordings, mice were habituated to being handled by the experimenter. Mice were restrained by hand while awake to quickly clean the surface of the GRIN lens with a triangular sponge (Sugi sponge points, Questalpha) soaked briefly in 70% ethanol and then connected to the head-mounted microscope.

### Recordings during free locomotion in the presence of visual landmarks

Mice were first screened for the presence of GCaMP6s-expressing and head-direction-tuned cells. The LED cylinder was set to display three distinct landmarks: i) a black rectangle spanning the full height of the cylinder and 45 degrees of visual angle, ii) a green triangle spanning the full height of the cylinder, with the base at the bottom spanning 45 degrees of visual angle, and iii) a purple circle with its center positioned at 48 cm from the base of the cylinder and diameter spanning 45 degrees of visual angle. The three landmarks were positioned equidistant to each other and were already displayed when the mouse was placed inside the cylinder. Mice were placed on a round, black, plastic platform (31.5 cm diameter, which was attached to the central enclosure housing the motor described above) in the center of the LED cylinder. The platform was cleaned thoroughly with 70% ethanol before and after each mouse. The excitation LED, gain, and electronic focus settings of the microscope were adjusted to maximize the signal quality and number of cells in focus. Mice in which no cells were visible after approximately 5 minutes of testing different imaging settings while the mouse was freely moving in the arena were not used for subsequent recordings. For mice in which cells were visible, a 20-minute recording was performed, in which the mice were freely moving around the platform and the behavior was recorded with a camera as described above. Immediately after this recording, the mouse was disconnected from the microscope and returned to its home-cage. The data from this recording was analyzed as described below, and mice with a sufficient number of cells tuned to head direction were used for subsequent recordings.

On a subsequent day, we performed a 13-minute-long recording under the same conditions as described above, immediately followed by another recording under head-fixation (during which the LED screen displayed a grating pattern, described in the following section), to confirm whether we can decode head direction during free locomotion and head-direction shifts during passive head rotations from the same cells, using the same decoder.

### Recordings during head-fixation

We performed multiple experiments in which transient visual and/or vestibular motion cues were presented to head-fixed animals. The mouse was head-fixed in the center of the cylindrical arena on a breadboard (Thor Labs) using a titanium head-bar. Throughout the experiment, the animal was standing on a sponge covered with tissue paper (with the rationale that the sponge is soft and therefore reduces motion artefacts caused by the mouse pushing down with its paws) and its body was covered and gently restrained with a black textile jacket to reduce motion of the body relative to the head. The breadboard, and with it the head-fixation apparatus, could be rotated by a brushless DC motor. The motor was controlled using the open-source solution ODrive (https://odriverobotics.com/). A control loop for motor position and velocity was running on the motor controller at a frequency of 8 KHz, using estimates of motor position and velocity from an encoder. We customized the firmware of the motor controller to allow for precise control of sinusoidal velocity trajectories ω(*t*) = ω_max_ sin(π*t*/*T*) with a stimulus duration of *T* = 3 s and a peak velocity ω_max_. Execution of these trajectories could be triggered in a Python script via a USB connection to the motor controller. In all experiments, trials were presented in random order, and the inter-trial interval was set to 2 s. We conducted two kinds of experiments in bright conditions and one kind of experiment in dark conditions. In the two kinds of experiments conducted in brightness, the cylindrical screen displayed a pattern of vertical black-and-white stripes, each of the 32 pairs of stripes covering 11.25 degrees of the visual field. The scene was controlled using PsychoPy^54^, and the velocity trajectories of the scene rotations were programmed to match those of the platform rotations.

### Passive rotations in brightness with stationary scene

In one set of experiments on wild-type animals, only the platform was rotated with a peak velocity ω_max_ that was varied between -180 deg/s (CW) and 180 deg/s (CCW) in steps of 45 deg/s while the scene remained stationary. Each stimulus was presented in 15 trials. The recording lasted 13 minutes (including pre- and post-stimulation periods) and was conducted immediately after the recording session in which the animal moved freely for 13 minutes where the LED screen displayed three visual landmarks. The data acquired during active and passive motion was preprocessed together. For all other head-fixed recordings (protocols described below), no freely moving recording was performed immediately prior to the head-fixed recording, and mice were directly head-fixed as described above and placed into the experiment arena.

### Recordings with all combinations of head and scene rotations

In another set of experiments, on both wild-type and *FRMD7^tm^* mice, the platform and the scene were rotated independently. We used the same velocity trajectory for both platform rotation and scene rotation with potentially differing peak velocities. Peak platform and scene velocities were independently varied between -180 deg/s and 180 deg/s in steps of 45 deg/s. This resulted in 81 combinations of peak platform and scene velocity. Each stimulus combination was presented 3 times per recording session, and we repeated the recording session three times on different days for each mouse to obtain sufficient statistics. Most recordings lasted 22 minutes with a stimulation duration of about 20 min and baseline periods at the beginning and end; for two wild-type mice and two *FRMD7^tm^*mice, 5 trials for each stimulus combination were presented instead of 3. We decided in favor of presenting only 3 trials in later experiments as we noticed a gradual decrease of the overall activity level over the course of individual recording sessions. For one *FRMD7^tm^* mouse, we were able to acquire only two recordings instead of three. In the experiments on the first three wild-type mice we did not track eye movement while for the remaining 7 wild-type mice we did track eye movement.

### Passive rotations in darkness

For both wild-type and *FRMD7^tm^* mice, we conducted passive rotations in darkness, with the same set of platform rotations (30 trials of each rotation speed, 15 CW and 15 CCW) as presented during the passive rotations in brightness, but without a prior phase of active locomotion. In dark conditions, the computer monitor as well as room lights were turned off. Windows were blacked-out and additional light sources within the room were coated by a black electrical tape and/or blackout fabric (BK5, Thor Labs). For the first three wild-type mice, the experiment in darkness was conducted twice on different days. For these three mice, two infrared LEDs (940 nm, Everlight Electronics) were attached to the microscope to enable verification of platform rotations from the top-view videos. For subsequent mice, the experiment was only conducted once as we found that the data from one experiment was sufficiently informative. For these mice, we acquired videos of the eye; thus an infrared light (850 nm, Thor Labs, with a longpass glass filter – cut-on: 830 nm, Schott RG830, Edmund Optics – attached in front of it) was present. However, the pupil was dilated to such an extent that pupil tracking was not possible. Nonetheless, the infrared light allowed verification of platform rotations via the top-view videos.

### Recordings with scene rotations during free locomotion

Following the same general recording procedure as before, for wild-type mice we conducted another recording during free locomotion with the scene sometimes transiently rotating during the experiment. In short, mice were restrained by hand and connected to the microscope. A red and a green LED were attached to the microscope to enable tracking of the head direction in the videos recorded with the top-view camera. Mice were placed on a round, black, plastic platform (31.5 cm diameter) in the center of the arena. The platform was thoroughly cleaned with 70% ethanol between each mouse. The LED cylinder displayed the same grating pattern as in the head-fixed recordings. This scene was stationary during most of the recording time. After a baseline period of 90 seconds to 5 minutes, 15 CW and 15 CCW scene rotations were presented in a random trial order and with random intertrial intervals of 30-60 seconds. The scene rotations had a peak velocity of either -45 deg/s (CW) or 45 deg/s (CCW). The recordings lasted 30 minutes.

### Escape-to-shelter spatial navigation paradigm

Wild-type mice without GRIN lens implants were used for the spatial navigation paradigm. The paradigm was performed in the experiment arena described above, but with a different custom-made circular platform. The scene displayed on the LED cylinder consisted of the same grating pattern used in the head-fixed recordings. The platform was, made from white PVC and had a diameter of 55 cm. A 1.3 cm hole in the center allowed it to be screwed to the breadboard connected to the motor. There were 16 holes evenly spread around the edge of the platform, each with a diameter of 6 cm and the outermost hole edge 2.5 cm away from the edge of the platform. All but one of these holes were sealed with transparent plexiglass. The remaining open hole served as the entrance to a shelter hidden under the platform. The shelter was an 8.5×11.5 cm plastic box attached to the underside of the platform. Some cage bedding from the home-cage of the mouse being tested was placed inside the shelter and onto the center of the platform. In addition, some white chocolate flakes or cereal were sprinkled onto the center of the platform, to encourage the mice to visit the center of the platform.

The paradigm consisted of three sessions conducted on separate days, each with one trial. The trial corresponded to one of the following conditions: control, platform-only rotation, or platform-and-scene rotation. Before any trial, a baseline period (>2 minutes) allowed mice to spontaneously find the shelter and habituate to entering and exiting the shelter. A trial was triggered when the mouse spontaneously exited the shelter and walked to approximately the center of the platform. In a control trial, once the mouse reached the center of the platform, an auditory stimulus was presented to elicit an escape response. In a platform-only rotation trial, once the mouse reached the center of the platform, the platform was rotated either CW or CCW at a peak speed of 45 deg/s (with the same trajectory as used in the head-fixed rotations). Immediately after the platform rotation finished, the auditory stimulus was presented. Platform-and-scene rotation trials only differed in that the scene was rotated in synchrony with the platform (with the same trajectories as the head-and-scene rotation during head-fixed experiments). In the first and second sessions, the trial corresponded to either a platform-only rotation or a platform-and-scene rotation, with the order counterbalanced across mice. In the third session the trial was a control trial for all mice. A different auditory escape stimulus was used on each day to minimize the risk of habituation. All stimuli were based on natural sounds. Specifically, the stimuli were eagle, hawk, and owl vocalizations. The stimuli were edited using the open-source software Audacity, such that each stimulus was composed of three repeating vocalizations (either eagle, hawk, or owl, respectively) totaling approximately 4.5 s. The stimuli were presented at 90-100 dB SPL measured at the center of the platform from a speaker (Pettersson L400) mounted above the platform and connected to a power amplifier (t.amp E-800, Thomann). Trials were triggered manually by the experimenter with a keyboard press using PsychoPy^54^ (which was also used to present the scene). The behavior of the mouse was recorded at a frame rate of 30 Hz with a camera (a2A2590-60ucPRO, Basler; wide angle lens #89-524, Edmund Optics) mounted centrally above the platform. Frames were timestamped by sending the camera exposure status to the microscope DAQ via BNC cables. Timestamp signals of the trials were also sent to the microscope DAQ via a BNC cable from a LabJack. In addition, a microphone (Ultragain Pro Mic 2200, Behringer) was placed near the speaker, and its output was also sent to the microscope DAQ via BNC cable as additional confirmation of the delivery and timing of the auditory stimulus.

The escape endpoint angle for each trial was quantified in Fiji^55^. A circle was drawn on the platform with the circle center matching the platform center and a radius such that the circle was tangential to the hole boundaries at their most interior points. The first frame after the escape stimulus presentation in which both ears of the mouse crossed the circle was taken as the endpoint of the escape response, and the center point of a line connecting the two ears was taken as the location of the escape endpoint. We then defined the escape angle as the angle between the vector from the platform center to the escape endpoint, and the vector from the platform center to the center of the shelter start position. To align CW and CCW trials, escape angles from CW trials were multiplied by -1. Thus, an escape angle of zero degrees refers to an escape to exactly the center of the shelter start position, and an escape angle of 85.9 degrees (the amount the platform and shelter were rotated by) refers to an escape exactly to the center of the shelter end position. Analysis was performed using MATLAB (MathWorks) and the CircStat toolbox for circular statistics^56^. 3 mice were excluded because of not entering the shelter during the baseline period before any trial presentation, and 2 mice were excluded because of a lack of an escape response in any direction.

### Histological verification of GRIN lens positions

After the completion of all recordings, mice were briefly anesthetized with isoflurane and given an overdose of Ketamine (120 mg/kg) and Xylazine (16 ml/kg) via an i.p. injection, and transcardially perfused with 4% PFA. The brain was removed and post-fixed in 4% PFA overnight at 4 °C and subsequently stored in PBS at 4 °C. Subsequently, 120 µm thick coronal sections were made with a vibratome (Leica VT1000S). Sections were immunostained with an anti-GFP primary antibody (1:1000, 600-101-215 goat anti-GFP, Rockland), followed by an anti-goat secondary antibody (1:500, Alexa Fluor 647 or 568 anti-goat, host: donkey, A21447 or A11057 Invitrogen) and Hoechst 33342 (Invitrogen). Sections were mounted on microscopy slides using ProLong Gold (Invitrogen). Fluorescence images of the sections were acquired with a spinning disc confocal microscope (Olympus IXplore SpinSR). Z-stack tile scans were acquired at 4X or 10X magnification (Olympus UPLXAPO 4X Objective or Olympus UPLXAPO 10X Objective, respectively). Stitching of tiles and maximum intensity projection was done using Olympus cellSens Dimension software. Image brightness and contrast was adjusted using Fiji^55^. For each mouse, the brain section which corresponded to the center of the implantation site (in the anterior-posterior dimension) was identified by inspection of the lesion made by the GRIN lens implant. To verify the position of the GRIN lens, these sections were manually overlaid with a reference image^57^ and the position of the GRIN lens tip was marked in Adobe Illustrator.

### Data preprocessing

Fluorescence videos were spatially downsampled by a factor of 2. Subsequently, we spatially bandpass filtered the frames with a low-frequency cutoff of 0.005 oscillations per pixel and a high-frequency cutoff of 0.500 oscillations per pixel. Finally, the videos were motion corrected. These steps were done using the proprietary Inscopix Data Processing Software.

Fluorescence traces of individual neurons were obtained using an open-source Python implementation of the CNMF-E algorithm^36,37^. First, a number of components were extracted from the video using the “corr pnr” initialization method based on pixel-specific local correlations and pixel-specific peak-to-noise ratios. Next, part of the extracted components were rejected based on thresholds for the signal-to-noise ratio (SNR) and the space correlation, whereby definitions of the quantities are as in^37^. If the SNR of a component was above 4.5, it was rejected if the space correlation was below 0.45. If the space correlation was above 0.7, the component was rejected if the SNR was below 3.5. If the SNR was below 4.5 and the space correlation was below 0.7, the component was also rejected. In addition, we rejected components that predominantly exhibited slow dynamics, which might be a sign of background activity or the apoptotic death of the neuron^58^. In detail, using a threshold frequency ω_thr_=0.05 Hz and a maximum frequency ω_max_=10 Hz, we defined a slow-to-fast ratio (SFR) as the energy of the temporal trace ƒ*(t)* in slow frequencies to the energy in fast frequencies:

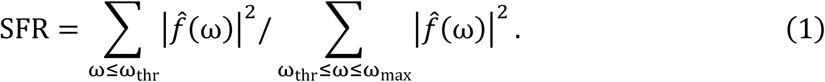

Here 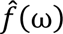 is the discrete Fourier transform. A component was rejected if the SFR exceeded a threshold value of 10. Finally, individual components were rejected based on experimenter judgement (e.g., because a neuron was not consistently in the field of view, the CNMF-E algorithm extracting the calcium traces erroneously split a single neuron into two spatial components, etc.).

Since the GCaMP6s sensor we used has slow dynamics, we deconvolved individual fluorescence traces. This was achieved with OASIS, an established activity extraction algorithm that estimates a first-order autoregressive model^59^. We chose a decay factor of γ=0.9 which corresponds to a half-life time of *t*_1/2_=-Δ*t* / log_2_ γ ≈ 0.55 s, where Δ*t*=1*/*12 s is the time between frames. This is close to the half-life time that has been reported for the GCaMP6s sensor in-vivo^60^. Due to the sparsity of the deconvolved traces, they were smoothed using a Gaussian filter with a standard deviation of 3 frames that was truncated at one standard deviation.

### Tracking of head direction and angular head velocity

Animal behavior was recorded at 30 frames per second. The head direction was tracked using a red and a green LED mounted to the left and the right side of the microscope on the head of mouse, respectively. For each individual frame, two binarized images were obtained by thresholding in HSV color space. After hole-filling and morphological opening operations on each binary image, the largest connected area was extracted. If the largest area consisted of least 100 pixels, the LED position was defined as the center of mass of this area. Otherwise, the LED tracking was considered unsuccessful, and the corresponding frame was excluded from analyses related to the head direction. Frames were processed with OpenCV^61^, and Ray^62^ for parallelization. The head direction was subsequently defined as the angle between the x-axis of the Cartesian coordinate system of the camera image and the vector connecting the red LED position to the green LED position plus 90 degrees. We obtained the angular head velocity by computing the difference quotient of the head direction as a function of time using pairs of consecutive frames, whereby frame-to-frame changes of the head direction were mapped to the interval from -180 deg to 180 deg. The head-direction trace was aligned to the calcium imaging video by linear interpolation. The interpolation was performed on intervals with successful LED tracking. For interpolation, the periodic head-direction trace was unwrapped; after interpolation, periodic boundary conditions were enforced again.

### Dimensionality reduction

By means of manifold learning, the high-dimensional population activity data was embedded into a two-dimensional embedding space to uncover the circular structure (“manifold”) of the data. The polar angle in the embedding space can be understood as the head direction encoded by the neuronal population activity.

In detail, the high-dimensional population activity data was mapped to a two-dimensional space using Laplacian Eigenmaps^34^, a nonlinear dimension reduction method. This method requires an affinity matrix that measures similarities between data points.

We computed a custom affinity matrix. First, for data obtained under head-fixed conditions, the data was trimmed to the interval starting 5 s before the first trial and ending 5 s after the last trial. In the case of recordings during free locomotion, the entire dataset was kept. To normalize the data, a quantile transform was applied to each smoothed and deconvolved calcium trace, mapping the data to a uniform distribution. Data points exactly zero were excluded from this normalization and consistently mapped to zero. The affinity matrix was obtained from the normalized high-dimensional population activity vectors.

The affinity between two population activity vectors *x* and *y* was defined by

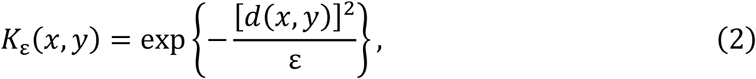

where ε is a global scale parameter and d(*x*,*y*) = 1 − ρ(*x*,*y*) is a dissimilarity measure based on the sample Pearson correlation ρ(*x*,*y*) between the vectors *x* and *y*. Since the sampling of directions might have been non-uniform, e.g., because the animal spent more time with its head pointing into a specific direction, the affinity was adjusted to

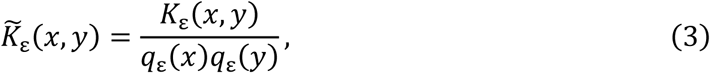

where *q*_ε_(*x*) = ∑_*y*_ *K*_ε_(*x*, *y*), as proposed in^63^. Finally, the affinity matrix was set to 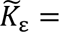 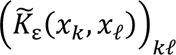 where *k* and *ℓ* denote points in time.

We used Laplacian Eigenmaps to map the activity vectors into a two-dimensional plane using the scikit-learn^64^ implementation of Laplacian Eigenmaps with symmetrically normalized Laplacian. In short, after removing self-loops from the affinity matrix by setting 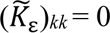, the symmetrically normalized Laplacian is given by 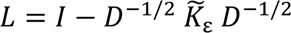, where *I* is the identity matrix and *D* is a diagonal matrix with entries 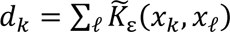.

The coordinates 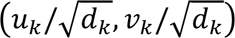 of the representation of the high dimensional population activity at time point *k* in the embedding plane were obtained from the eigenvectors *u* and *v* corresponding to the second and third smallest eigenvalues of *L*. The eigenvector with the smallest eigenvalue (= 0), corresponding to the (unique) connected component of the graph defined by the affinity matrix (understood as an adjacency matrix) was dropped as common in Laplacian Eigenmaps.

We decoded the head direction by mapping each high-dimensional population activity vector to the polar angle of the data point representing the population activity in the two-dimensional embedding space. However, if appropriate we changed the orientation of the decoded angle and potentially applied a global rotation. Before describing this adjustment in detail, we explain how the global scale parameter *ε* used to compute the affinities was chosen.

The mapping depends on the value of the global scale parameter ε. The precise value of ε was determined separately for each recording. First, embeddings were computed for values of *ε* between 0.01 and 0.20 in steps of 0.005. For each embedding, the Cartesian coordinates in the two-dimensional embedding plane were transformed to polar coordinates. We expected the empirical distribution of polar angles to not strongly deviate from a uniform distribution on [−π, π]. Therefore, if the dissimilarity between the empirical distribution and a discrete uniform distribution, as quantified by the Kullback-Leibler divergence, exceeded the threshold of 0.4, the corresponding value of *ε* was not considered further. Here, the empirical distribution of polar angles was obtained by binning the data into 36 evenly spaced intervals.

Among the remaining values of ε, the value was chosen for which the embedded data deviated the least from a one-dimensional structure. The deviation was measured by the median (across data points labeled by *k*) of the absolute value of *r_k_* / *r*(ϕ*_k_*) − 1 where (*r_k_*, ϕ*_k_*) are the polar coordinates of data point *k* and *r*(φ) is a curve fit. The curve fit *r*(*φ*) is the weighted median of *r_k_* with weights *w*_*k*_ = exp {−[φ_*k*_ − φ + π (mod 2π) − π)]^2^}/σ^2^}. The scale parameter σ was set to 0.05 rad. Finally, values of ε were not considered if the origin (0,0) was not in the convex hull of the fitted curve.

The orientation of the decoded head direction was adjusted differently between freely moving and head-fixed recordings with manipulations of the visual scene. In the former case, we split the temporal traces of the actual and the decoded head direction into chunks of 30 s. In a chunk of this duration, we expected a high positive correlation between the head direction and the decoded head direction even if the two variables would drift away from each other on a slower time scale. For each chunk, the circular correlation^65^

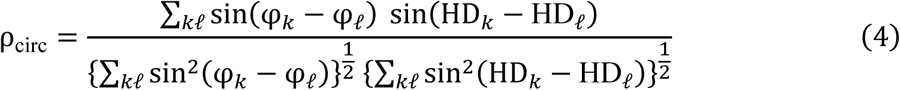

between embedding angle φ*_k_* and head direction HD*_k_* was computed. Here, *k* and *ℓ* label time points. If the mean circular correlation across chunks was negative, we reversed the orientation of the embedding angle. Otherwise, the orientation remained unchanged.

For head-fixed recordings, we adjusted the orientation of the decoded head direction based on trials with “natural” stimulation where only the platform rotated but not the scene. We chose the orientation which maximized the Pearson correlation between the decoded head-direction shift —as specified in the following section —and the peak platform velocity.

In the absence of visual landmarks, the decoded head direction was taken as identical to the — potentially reversed —embedding angle. In the presence of visual landmarks, we subtracted *c* = arg ∑_*k*_ *e*^*i*(φ*k*−HD^_*k*_^)^ (with the average across frames with successful LED tracking) from the decoded head-direction trace. This minimized *f*(*c*) = ∑_*k*_ d(φ_*k*_ − *c*, HD*_k_*) with the circular distance d(α, β) = 1 − cos(α − β). However, we quantified the decoding error as the median absolute deviation as this measure is more intuitively understood than the mean circular distance. In calculating the absolute deviation, we took into account periodic boundary conditions, that is, the decoding error is given by the median of |φ*_k_* – *c* − HD*_k_* + *π* (mod 2*π*) − *π*|.

### Decoded head-direction shift

We quantified the total shift Δφ of the decoded head direction (“decoded head-direction shift”) in response to transient stimulation by unwrapping the decoded head direction *φ* and subtracting the mean decoded head direction φ_before_ across a 1 s interval immediately before stimulus onset from the mean decoded head direction φ_after_ across a 1 s interval immediately after stimulus offset. The time series were obtained by subtracting φ_before_ from the unwrapped decoded head-direction trace.

We defined the normalized shift in percent differently for experimental conditions in which the platform rotates and conditions where only the scene rotates. In the former case, the normalized shift corresponded to the ratio between the decoded head-direction shift and the actual head-direction shift. In the latter case, it corresponded to the ratio of the decoded head-direction shift and the angular shift of the scene. For each of the experimental conditions involving head rotations, data from CW head rotations was sign-inverted to align with data from CCW head rotations. Data from CCW scene rotations was inverted to align with data from CW scene rotations.

### Cross-correlation analysis between decoded head-direction trace and actual head-direction trace

In the deconvolution algorithm, we only took into account the decay of the fluorescence signal, but not the initial rise of the signal. The rise time of GCaMP6s as quantified by the in-vivo time-to-peak is around 200 ms^60^. Therefore, for the analysis of the decoded head direction measured in response to visual-vestibular stimulation, we corrected for a potential systematic lag of the decoded head direction.

To correct for a potential lag of the decoded head direction, we quantified the delay that maximizes the correlation between the decoded head direction and the actual head direction during active locomotion. The activity of head-direction cells in the anterior thalamus has been found to anticipate head direction by 20 ms to 40 ms on average in rats^66–69^. As we recorded at a frame rate of 12 fps corresponding to a frame duration of ≈ 83 ms, using the lag with maximum correlation was adequate. Note however that, not invalidating the calibration approach, anticipatory time intervals have been found to be increased during passive motion^70^. For calibration, we used the first part of the recording session combining active and passive motion, i.e., the part in which the animal is freely locomoting in the presence of visual landmarks. For all animals, the correlation was maximized by advancing the decoded head-direction trace by either one or two frames. On average across animals, this corresponded to a delay of 100 ms of the decoded head direction with respect to the actual head direction. In the analysis of the decoded head-direction shift during passive motion, we advanced the decoded head-direction trace by this amount.

### Head-direction tuning

Tuning curves with respect to the head direction were obtained by computing the average activity in each of 36 equally spaced angular bins. We defined a neuron’s preferred head direction as the center of the bin with the highest average activity. The strength of directional tuning was quantified using the Rayleigh vector length which is the resultant length of a sample of vectors *p*_*k*_*e*^*i*α^_*k*_ where *p*_*k*_is the ratio of the mean activity measured in bin *k* and the sum of the mean activities across bins, and α_*k*_ is the center of bin number *k* in radians. We considered a neuron “tuned” to head direction if the measured Rayleigh vector length exceeded the 99th percentile of a sample of randomized vector lengths (see Statistics). Note that for the dimension reduction all neurons were included, independently from the significance testing.

### Extraction of pupil position

We measured the eye-in-head position in degrees following a methodology similar to^51^. Using DeepLabCut^71^, we trained a deep neural network to track the pixel locations of nasal and temporal eye corner as well as temporal and nasal pupil edge. We considered tracking as successful if the likelihood of each of the four points (as quantified by DeepLabCut) exceeded 95%. This was the case for between 97.4% and 99.5% of frames for individual animals. One network was trained for each animal based on a manually labeled dataset. Initially, we labeled 20 frames for each of the three passive rotation experiments in which platform and scene velocity were independently controlled, and we optimized the network on this training dataset, consisting of a total of 60 frames, for 100,000 iterations. If the tracking was not yet satisfactory after this training phase, we added additional frames to the training dataset and continued training.

As we were interested in rotations around the yaw axis, we projected the pixel locations of the tracking points onto the horizontal axis. Pixel values were converted to mm assuming that the average distance between nasal and temporal eye corner (across frames with successful tracking) is approximately 3.2 mm. This value of this distance was used in^51^, and we verified the adequacy of this value.

We calculated the absolute pupil position as the average of nasal and temporal pupil edge. Subsequently, the pupil position was converted to an eye-centered coordinate system and then to an angular estimate with a small-angle approximation using an effective eyeball radius of 1.25 mm^72^.

To account for potential, small horizontal movements of the whole eye, the eye center was not simply taken as the mean of nasal and temporal eye corner. Instead, the eye center was defined as the temporal eye corner (which in the image was to the right of the pupil) minus the distance between nasal and temporal eye corner multiplied by 1 minus the average relative pupil position. Here, the relative pupil position is the ratio between the absolute pupil position minus the nasal eye corner (which in the image was to the left of the pupil) and the distance between nasal and temporal eye corner.

Given the temporal trace of the pupil position in degrees, linear interpolation was performed for isolated frames of unsuccessful tracking. The temporal trace was subsequently filtered using a Gaussian window with a length of 5 frames (≈ 83 ms) with a standard deviation of 1 frame that was truncated at 2 frames. Finally, we obtained the pupil velocity over time by taking the difference quotient of the pupil position trace with a step of 1 frame. We refer to the pupil velocity as the eye velocity.

### Estimation of slow-phase eye velocity and angular eye shift

We were interested only in the slow-phase dynamics of eye movement. Therefore, we detected quick phases and excluded these from subsequent analyses. The detection was performed at the level of individual trials. Trials with frames of unsuccessful eye tracking (after interpolation of isolated frames of unsuccessful eye tracking) were excluded from the analysis of slow-phase eye movement. For each trial, we first determined the average direction of eye movement. The reason for this is that the slow-phase eye movements typically occurred in a dominant direction, and most time points corresponded to a slow phase. We then first found quick phases in which the eye moved in the direction opposite to the mean direction of slow-phase eye movement. This was done by detecting intervals in which the eye speed consistently exceeded a threshold of 10 deg/s. The threshold value was set so low to include the beginning and the end of the quick phase. The subset of intervals in which the peak eye speed exceeded a threshold of 150 deg/s and that lasted at most 12 frames (= 200 ms) were classified as quick phases. Next, we extracted occasional quick phases that occurred in the same direction as the mean direction of slow-phase eye movement using the same detection thresholds. However, detected intervals that occurred less than 100 ms before or after one of the previously identified quick phases were not classified as quick phases. We found that this methodology captures most quick phases.

After removing quick phases, a polynomial of degree 4 was fit to the eye velocity by means of a Huber regression which is a linear regression that is robust to outliers. In the regression, time points were weighted according to the inverse of a kernel density estimate obtained with a Gaussian kernel with a standard deviation of 150 ms. This ensured that periods in which lots of quick phases occurred are still adequately taken into account. The fit yielded an estimate of the slow-phase eye velocity over time, and interpolated periods of quick phases, i.e., predicts the would-be eye velocity if there was no need for quick phases to recenter the eye.

Finally, we obtained the total angular eye shift during the stimulation as the temporal integral of the slow-phase eye velocity as estimated by the fitted polynomial function. We defined the “eye turn” as the additive inverse of the resulting quantity; the additive inverse was used such that the sign of the eye turn matched that of the decoded head-direction shift. Analogous to the decoded head-direction shift, the normalized eye turn was calculated by dividing the eye turn in degrees by the angular shift of the head (if the head rotated) respectively scene (if only the scene rotated).

### Model

We developed a joint model of visual-vestibular interaction for both reflexive eye movements and angular path integration in the head-direction system. The interaction happens between (i) a visual signal and (ii) a vestibular signal, which are evoked by image motion on the retina and head motion, respectively. We modeled the interaction between visual and vestibular signals by (iii) a differential equation describing the dynamics of a multisensory head velocity estimate. The head-direction system integrates this velocity estimate over time, (iv), to yield a current estimate of head direction. The velocity estimate also drives reflexive eye movements, (v), that typically go in the direction opposite to the estimated head rotation. However, the eye movements have a temporal lag due to the mechanical properties of the oculomotor plant, and we modeled slight differences between the decoded head-direction shift and the sign-inverted angular eye shift by allowing the vestibular signal to have a direct influence on the eye velocity instead of only an indirect influence via the model’s head velocity estimate. Finally, we describe, (vi), how the numerical values of the model parameters were obtained.

#### (i) Visual signal

Rotation of the head around the yaw axis, (artificial) rotations of the scene around the head as well as eye movements cause whole-field rotational image motion on the retina. The rotational component of the retinal image slip (“retinal slip”) is given by

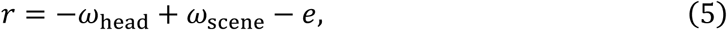

where *ω*_head_ is the angular head velocity, *ω*_scene_ is the angular velocity of the scene and *e* is the eye velocity. Retinal slip generates a visual signal

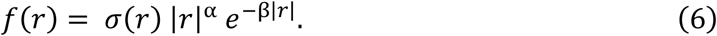

Here, *σ* is the sign function and α and β are parameters specified later. The use of a nonlinear transform of the retinal slip is in line with classical models of the optokinetic reflex^73,74^. The nonlinearity presumably arises from response characteristics of direction-selective ganglion cells (DSGC) in the retina. In mice, responses of two subtypes of On DSGCs and On-Off DSGCs likely involved in driving the optokinetic reflex steadily decrease for image speeds larger than 5 deg/s^41^. Although there are differences in the response characteristics between subtypes of DSGC, we restrict ourselves to a single visual channel. The functional form *f*(*r*) we use captures the main response characteristics of these direction-selective retinal ganglion cells, and has been used before to model optokinetic nystagmus^75^.

#### (ii) Vestibular signal

In mammals, rotational head motion around the yaw axis is detected in the vestibular organ by the semicircular canals^76^. Responses of neurons innervating the semicircular canals are well described by a linear dynamical system^77^. Specifically in mice, a study^40^ found that vestibular afferents classified as regular or irregular are both well described by a linear dynamical system. Based on this study, and neglecting a fast time constant *<*25 ms which has no material impact for the trajectories used in the present study, we obtained the vestibular signal *c* from the differential equation

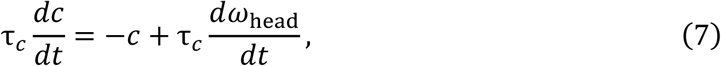

where 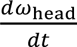 is the angular head acceleration, and the time constant τ*_c_* = 3 s is set to a value between the values of the (slow) time constant reported for regular afferents and irregular afferents. This means the vestibular signal corresponds to an exponentially filtered version of angular head acceleration.

#### (iii) Differential equation for visual-vestibular interaction

We model the dynamics of an estimate 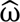 of angular head velocity which combines visual with vestibular information. This velocity estimate drives both compensatory eye movements and updates the current estimate of head direction in the head-direction system. The dynamics of 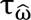 is described by a first-order differential equation:

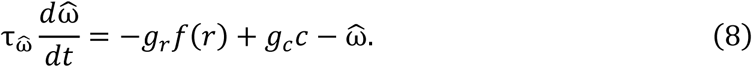

Here, 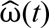 is the time constant of the multisensory estimation process, *g_r_* is the gain of the visual signal *f*(*r*) and *g_c_* is the gain of the vestibular signal *c*.

The visual gain *g_r_* = *g_r_*_0_ + γ*_r_* |*c*| has a static component *g_r_*_0_ but in addition we allowed that it increases with larger vestibular signal with a slope γ*_r_*. Moreover, the vestibular signal is suppressed at small values by multiplication with a gain *g_c_* = min{*g_c_*_0_ + γ*_c_* |*c*|,1} that is smaller than one for small vestibular signal and leaves the vestibular signal unchanged for large vestibular signal. For *g_c_*<1, the vestibular gain has a static component *g_c_*_0_ and linearly increases with a slope γ*_c_* in dependence on the magnitude of the canal signal, analogous to the visual gain. We allowed for a non-static vestibular gain based on findings of a velocity-dependence of the VOR gain^38^ and, for the sake of simplicity, we modeled the dependence as a saturating linear function. Since the vestibular signal generated in the inner ear has been reported to be linear^40^, we allowed the raw, linear signal to drive motoneuron activity and thereby influence the dynamics of the eye plant, see (v).

#### (iv) Integration of the velocity estimate by the head-direction system

We assume that the direction encoded by head-direction cells in the anterodorsal thalamus is the integral of the angular velocity estimate 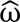(*t*) without any (positive or negative) temporal lag. Importantly, we do not model the integration process: there are numerous proposals how the integration process could be implemented in the head-direction system^20,21,78,79^. Further, the assumption that there is no temporal lag is a simplification: Head-direction cells in the rat AD slightly anticipate head direction during free locomotion^66,67^. Note also that in in rat lateral mammillary nucleus, which is upstream to the AD in the head-direction system^32^, head-direction cells anticipate head direction more than in AD, by about 40 ms to 70 ms^68,80^.This anticipation could, for example, be explained by adding an acceleration component to the velocity signal. We neglected these details due to the slow dynamics of the calcium indicator used in the current study.

#### (v) Dependence of eye movements on the velocity estimate and the vestibular signal

We hypothesized that reflexive gaze-stabilizing eye movements are driven by the same estimate of angular head velocity used by the head-direction system. Moreover, we allowed vestibular signals to directly influence the eye velocity in addition to the indirect influence via the velocity estimate used by the head-direction system. Specifically, we obtained the time evolution of the velocity of the eye, *e*, from the equation

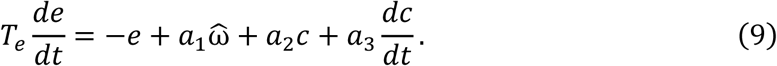

The eye velocity thus relaxes to a linear combination of the angular velocity estimate 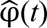, the vestibular signal *c*, and its temporal derivative, with a time constant *T_e_*, in line with the rather sluggish response of the oculomotor plant to due to its biomechanical properties^81^. The coefficients *a*_1_, *a*_2_, and *a*_3_ determine the relative influence of the three signals. The numerical values of these coefficients and the time constant *T_e_* were calculated from the regression analysis described below.

Integrating both sides of Eq. (9) results in the differential equation

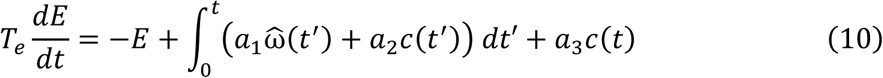

for the angular eye shift

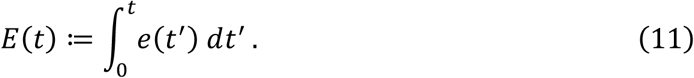

In practice, we dealt with the discretized version of Eq. (9),

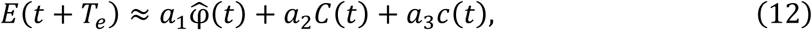

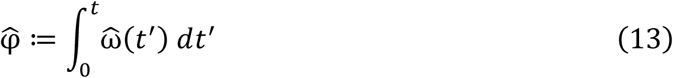

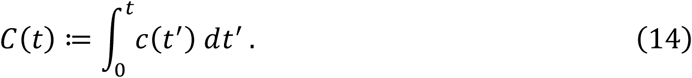

To obtain the coefficients *a*_1_, *a*_2_ and *a*_3_, we regressed the lagged, average, slow-phase eye shift, *E*(*t* + *T_e_*) on the average angular shift 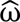 of the decoded head direction, the theoretical vestibular signal, and its integral. Specifically, we minimized the quadratic error given by

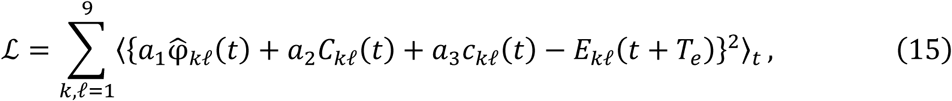

where *k* and *ℓ* index the peak platform velocity and the peak visual scene velocity, both ranging from −180 deg/s to 180 deg/s in steps of 45 deg/s, respectively. Before averaging the angular shift of the decoded head direction across trials and animals, we aligned the data to the same grid of time points by linear interpolation. The grid covered the period 0 s to 3 s (i.e., the stimulus duration), sampled at a frequency of *f* = 12 Hz. We systematically varied the lag *T_e_* of the oculomotor plant between 0 and 6 time points, i.e., between 0 s and 0.5 s, and determined the optimal value. The last *f* · *T_e_* time points were excluded from the regression, as the slow-phase eye shift was quantified only during the transient stimulation. To compare the same values of the slow-phase eye shift between the different lag values, the first 6 − *f* · *T_e_* time points were also excluded. The regression analysis resulted in an optimal lag of 2 time steps (≈167 ms) and in the parameter values *a*_1_ = −0.95, *a*_2_ = −0.17, and *a*_3_ = −0.18.

The time constant of 167 ms is at the lower end of the (frequency-dependent) time constant reported for mice in Ref.^44^, ranging between 130 ms and 450 ms. In particular, Ref.^44^ reported a dominant time constant of 350 ms at a frequency of 0.2 Hz, which is close to the frequency of 1/6 Hz used in our experiments. However, the study investigated responses to sustained stimulation; therefore, transient effects relevant to our case might not have been captured.

Since Eq. (10) has the same form as that of a model of the oculomotor plant^81^, the quantity

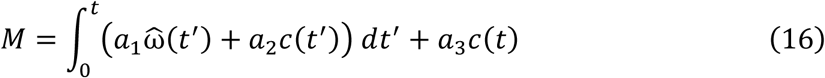

can be interpreted as the oculomotor command signal, encoded in the firing rates of motoneurons: the temporal integration (of a weighted sum of the estimated angular head velocity, 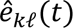, and the vestibular signal, *c*) can be performed by the oculomotor integrator. In a more detailed model, however, the oculomotor integrator might be leaky, to prevent noise from accumulating.

#### (vi) Estimation of model parameters

The model parameters α, β related to the visual nonlinearity, the time constant 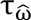, the visual gain parameters *g_r_*_0_, γ*_r_* and the vestibular gain parameters *g_c_*_0_, γ*_c_* were obtained by minimizing the mean squared error between the model prediction for the eye velocity and the measured eye velocity.

In detail, we minimized the weighted mean squared error

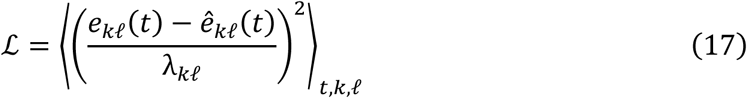

between the predicted eye velocity *e*^_*k*ℓ_(*t*) and the measured eye velocity *e*_*k*ℓ_(*t*) across time and the 45 experimental conditions (where data from CW and CCW platform rotations was aligned and aggregated) indexed by 1 ≤ *k,ℓ* ≤ 9, corresponding to combinations of peak platform velocity and peak scene velocity, using the BFGS (Broyden–Fletcher–Goldfarb–Shanno) algorithm. The experimental conditions were weighted by λ*_kℓ_* = λ_min_ + max*_t_* |*e_kℓ_*(*t*)| with λ_min_ = 10 deg/s to prevent a bias towards conditions in which the eye moves at high speeds. The fitted model parameters are listed in the following table; in the simulations of the model, the unit for angles was radians, and the unit for time was seconds.

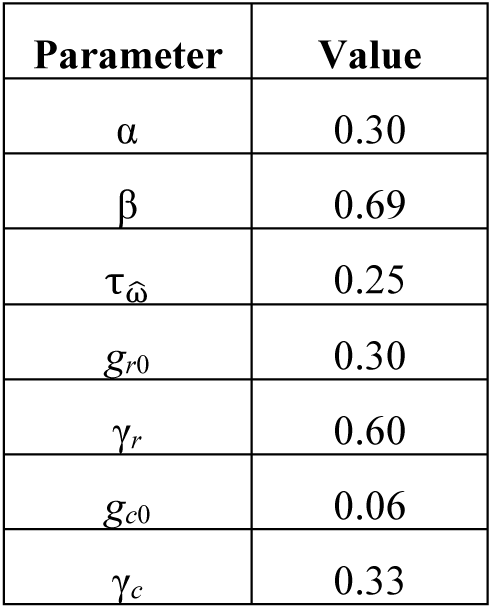

| Parameter | Value |
| --- | --- |
| $\alpha$ | 0.30 |
| $\beta$ | 0.69 |
| $\tau_{\hat{\omega}}$ | 0.25 |
| $g_{r0}$ | 0.30 |
| $\gamma_r$ | 0.60 |
| $g_{c0}$ | 0.06 |
| $\gamma_c$ | 0.33 |

### Bayesian model

Here, we describe the Bayesian model in detail. We consider a discrete-time hidden Markov model in which the angular head velocity ωω_*t*_ follows an unobserved Markov process. Transitions from one time point to the next are modeled as Gaussian:

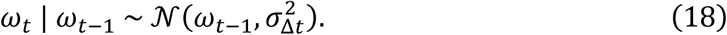

At each time point, there are three observations corresponding to a noisy visual signal *zₜ*, a noisy vestibular signal *c*_*t*_, and a noisy motor signal *m*_*t*_. For convenience, we aggregate these observations into the vector *x*_*t*_ = (*z*_*t*_, *c*_*t*_, *m*_*t*_).

First, the visual signal *z*_*t*_ is distributed normally with a mean obtained from a nonlinear transformation *f* of the retinal slip *r*_*t*_ = −ω_*t*_ − *e*_*t*_ (in the natural situation where the visual scene is stationary) and the standard deviation σ_*z*_:

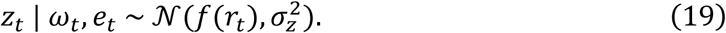

Here, the nonlinear transformation is given by the function

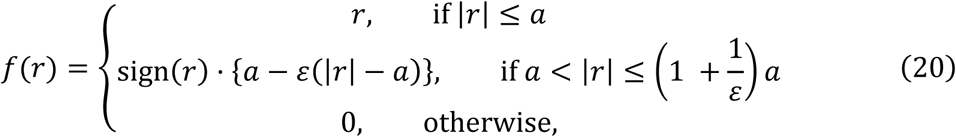

with parameters ε and *a* specified later on. As in the previously introduced mechanistic model of visual–vestibular interactions, the visual signal increases with retinal slip for small positive values, but decreases as the slip becomes larger.

Second, the nosiy vestibular signal arises from sensory transduction in the semicircular canals. For simplicity, we approximate the observation model by assuming that the vestibular signal has a normal distribution with mean identical to ωω*ₜ* and a standard deviation σσ_*c*_.

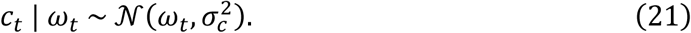

Nonetheless, when simulating observations (see further below), we take into account the dynamics of the semicircular canals as described by Eq. (7).

Third, the motor signal is normally distributed with a mean identical to and a standard deviation that depends on the angular head velocity:

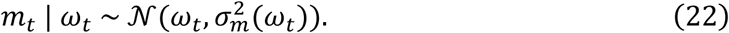

The standard deviation

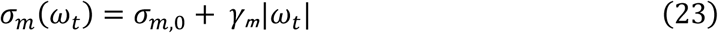

consists of a constant component σ_*m*,0_ and a component that increases linearly, with a slope *χ*_*m*,0_ with the magnitude of the angular head velocity. The latter is supported by experimental evidence for signal-dependent noise in the in the execution of movements^82–84^.

We would like to compute the posterior distribution *p*ω_*t*_| *x*_1_, …, *x*_*t*_) over the angular head velocity at time point *t* given all evidence up to that time point. This computation can be performed sequentially using a Bayes filter. First, we note that

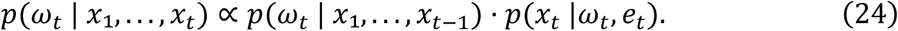

The predictive distribution *p*(ω_*t*_ | *x*_1_, …, *x*_*t*−1_) is given by

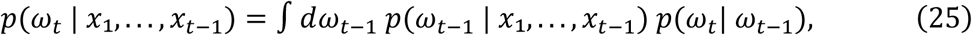

represting a convolution as the transition probability only depends on the difference between ω_*t*_ and ω_*t*−1_. To update the posterior distribution, the predictive probability is multiplied with the likelihood of the ‘observed’ sensory signals. Assuming conditional independence of the sensory modalities, the likelihood factorizes:

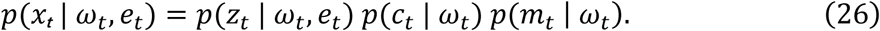

As an essential element of our probabilistic framework, we set the eye velocity equal to the additive inverse of the posterior mean at the previous time step:

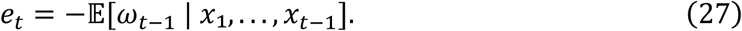

We implemented the Bayes filter to numerically compute the posterior distribution over time for each of our experimentally applied stimuli. The visual signal was set to

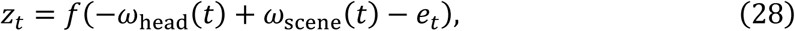

with the head and scene velocity profiles ω_head_(*t*) and ω_scene_(*t*) as used in the experiments. The vestibular signal was obtained from Eq. (7) and the motor signal was set equal to the zero. In the simulations, we set the time increment to Δ*t* = 0.025 s and the parameters determining the signal and noise structure to the values given in the following table.

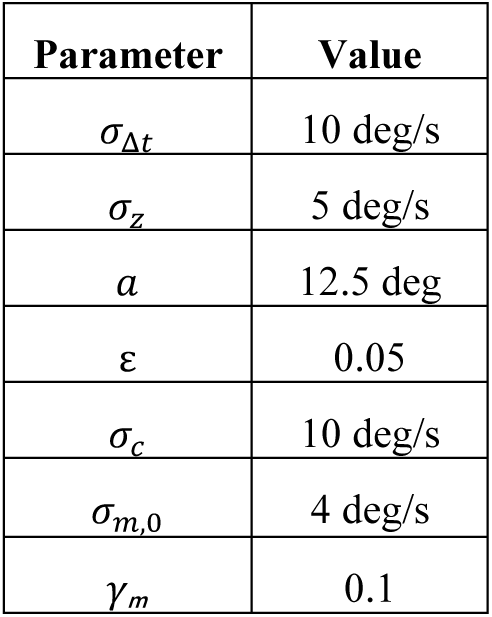

| Parameter | Value |
| --- | --- |
| $\sigma_{\Delta t}$ | 10 deg/s |
| $\sigma_z$ | 5 deg/s |
| $a$ | 12.5 deg |
| $\varepsilon$ | 0.05 |
| $\sigma_c$ | 10 deg/s |
| $\sigma_{m,0}$ | 4 deg/s |
| $\gamma_m$ | 0.1 |

### Statistics

We considered a neuron “tuned” to head direction if the measured Rayleigh vector length exceeded the 99th percentile of a sample of randomized vector lengths (Extended Data Fig. 1f). The randomized vector lengths were obtained by systematically advancing the direction trace between 5 s and 60 s in time, wrapping the end of the session to the beginning, and computing corresponding, randomized tuning curves. Tuning curves were computed by binning the calcium activity into n=36 equally spaced bins for each of the 658 neurons pooled across 10 recordings (one per animal).

For each animal, we tested the null hypothesis that the preferred head directions of different neurons spread uniformly over all angles using a Rayleigh test. The test was applied to all extracted neurons in each of the 10 recordings (one recording per animal, with n=40, 88, 36, 49, 48, 66, 97, 107, 89, 38 neurons, respectively) and yielded the following p-value per animal: 0.98, 0.33, 0.76, 0.87, 0.19, 0.45, 0.59, 0.28, 0.37, 0.33. Rayleigh’s statistic took the values: 0.02, 1.10, 0.27, 0.14, 1.69, 0.81, 0.52, 1.28, 0.99, 1.11.

We performed a paired t-test compare the animal-averaged decoded head-direction shift between head-only rotations and head-and-scene rotations in both (n=10) wild-type and (n=4). *FRMD7^tm^* animals (with 9 respectively 3 degrees of freedom). It was one-sided for wild-type animals to test the null hypothesis that head-synchronous visual-scene rotation does not reduce the decoded head-direction shift against the alternative that it does. It was two-sided for *FRMD7^tm^* animals to test the null hypothesis that the visual-scene rotation does not modulate the decoded head-direction shift against the alternative that it does. A Bonferroni-Holm correction was applied for testing four hypotheses (corresponding to the different peak stimulus speeds). For wild-type animals the t-statistics, ordered with increasing head speed, took the values -14.0, -22.2, -39.3, -14.2 and after Bonferroni-Holm correction, the p-values took the values 1.8×10^-7^, 5.5×10^-9^, 4.5×10^-11^, 1.8×10^-7^. For *FRMD7^tm^*animals the t-statistics took the values 1.11, -0.33, -1.12, -0.70 and the corrected p-values all took the value 1.0.

For both (n=10) wild-type and (n=4) *FRMD7^tm^* animals, one-sided t-tests were conducted (with 9 respectively 3 degrees of freedom) for the null hypothesis that the animal-averaged decoded head-direction shift evoked by scene-only rotation had mean zero against the alternative of a positive mean. The t-statistics and the Bonferroni-Holm-corrected p-values are listed in the table below.

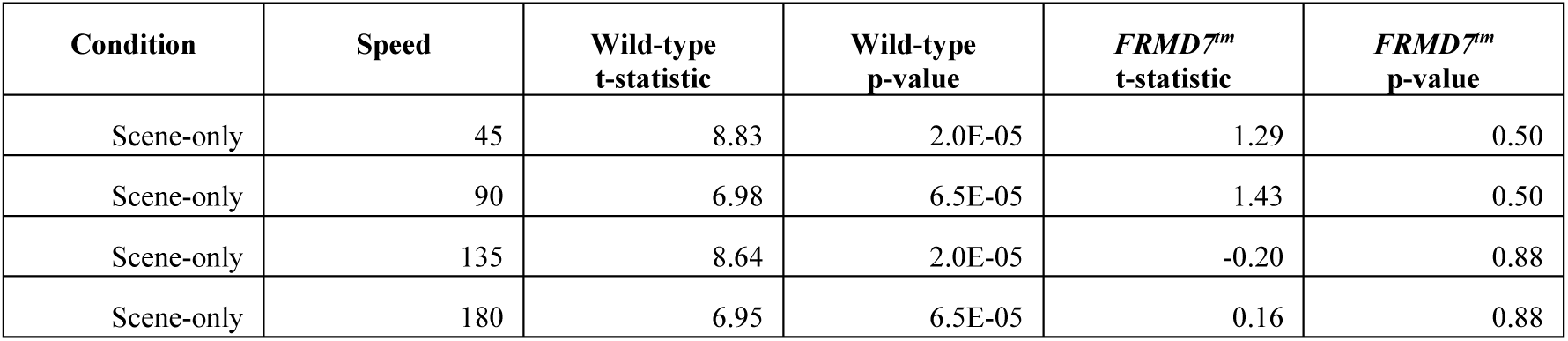

We tested whether the decoded head-direction shift in percent depended on the peak stimulus speed for head-only, head-and-scene as well as scene-only rotations (Fig. 1j). For each of the three conditions, a linear curve was fit to the animal-averaged data using ordinary least squares. For eight animals we repeated measurements 18 times (corresponding to 9 trials each for CW and CCW rotation) and for the remaining two animals we repeated measurements 30 times (corresponding to 15 trials each for CW and CCW rotation). Separate two-sided t-tests (on n=4 speeds×10 animals=40 samples; 38 degrees of freedom) were conducted on the slope parameter of the fitted curves. The t-statistics took the values 2.33 for head-only rotation, 9.21 for head-and-scene rotation and -8.53 for scene-only rotation. Bonferroni-Holm correction for testing three hypotheses was applied, yielding corrected p-values of 0.025 for head-only rotation, 9.5×10^-11^ for head-and-scene rotation and 4.6×10^-10^ for scene-only rotation.

To test whether the decoded head-direction shift was smaller in conditions in which the visual scene rotated compared to the head-only rotation condition, we performed one-sided paired t-tests for (n=10) wild-type (with 9 degrees of freedom). For each cohort and each peak head speed, we applied a Bonferroni-Holm correction to the 8 tests (4 for positive and 4 for negative scene velocities). The t-statistics and the corrected p-values are listed in the table below.

| Head speed [deg/s] | 45 |  | 90 |  | 135 |  | 180 |  |
| --- | --- | --- | --- | --- | --- | --- | --- | --- |
| Scene velocity [deg/s] | t-statistic | p-value | t-statistic | p-value | t-statistic | p-value | t-statistic | p-value |
| -180 | -4.80 | 2.0E-03 | -7.08 | 1.2E-04 | -5.94 | 3.3E-04 | -3.88 | 7.4E-03 |
| -135 | -4.47 | 2.3E-03 | -5.16 | 8.9E-04 | -6.75 | 1.7E-04 | -3.36 | 0.01 |
| -90 | -0.62 | 0.55 | -2.57 | 0.03 | -3.40 | 7.9E-03 | -2.40 | 0.04 |
| -45 | 2.67 | 0.99 | -1.22 | 0.13 | -0.29 | 0.39 | 0.41 | 0.65 |
| 45 | -14.05 | 5.0E-07 | -20.06 | 2.2E-08 | -17.95 | 5.9E-08 | -6.58 | 2.5E-04 |
| 90 | -16.42 | 1.9E-07 | -22.16 | 1.3E-08 | -28.92 | 1.2E-09 | -13.30 | 9.6E-07 |
| 135 | -14.68 | 4.1E-07 | -20.50 | 2.3E-08 | -39.28 | 8.9E-11 | -21.99 | 1.6E-08 |
| 180 | -16.53 | 1.9E-07 | -25.08 | 4.9E-09 | -28.84 | 1.2E-09 | -14.22 | 6.3E-07 |

We performed analogous, but two-sided, paired t-tests for the (n=4) *FRMD7^tm^* (with 3 degrees of freedom). The t-statistics and the corrected p-values are listed in the table below.

| Head speed [deg/s] | 45 |  | 90 |  | 135 |  | 180 |  |
| --- | --- | --- | --- | --- | --- | --- | --- | --- |
| Scene velocity [deg/s] | t-statistic | p-value | t-statistic | p-value | t-statistic | p-value | t-statistic | p-value |
| -180 | 1.01 | 1.00 | -0.07 | 1.00 | -2.23 | 0.67 | -0.16 | 1.00 |
| -135 | 2.20 | 0.93 | 1.53 | 1.00 | -2.74 | 0.50 | -0.26 | 1.00 |
| -90 | 1.94 | 1.00 | -0.15 | 1.00 | 1.52 | 1.00 | -0.12 | 1.00 |
| -45 | 0.07 | 1.00 | 1.06 | 1.00 | 1.42 | 1.00 | -0.93 | 1.00 |
| 45 | 1.11 | 1.00 | 1.12 | 1.00 | 0.33 | 1.00 | 0.12 | 1.00 |
| 90 | 0.24 | 1.00 | -0.33 | 1.00 | -4.37 | 0.18 | 0.65 | 1.00 |
| 135 | 0.24 | 1.00 | 2.05 | 1.00 | -1.12 | 1.00 | -1.26 | 1.00 |
| 180 | 0.97 | 1.00 | -0.32 | 1.00 | -1.47 | 1.00 | -0.70 | 1.00 |

For the (n=10) wild-type animals and conditions in which the scene velocity exceeded the head speed, we tested the null hypothesis the animal-averaged decoded head-direction shift had a population mean of zero versus the alternative hypothesis that the population mean is negative using a one-sided t-test (with 9 degrees of freedom). Bonferroni-Holm correction was applied. The t-statistics and the corrected p-values are listed below.

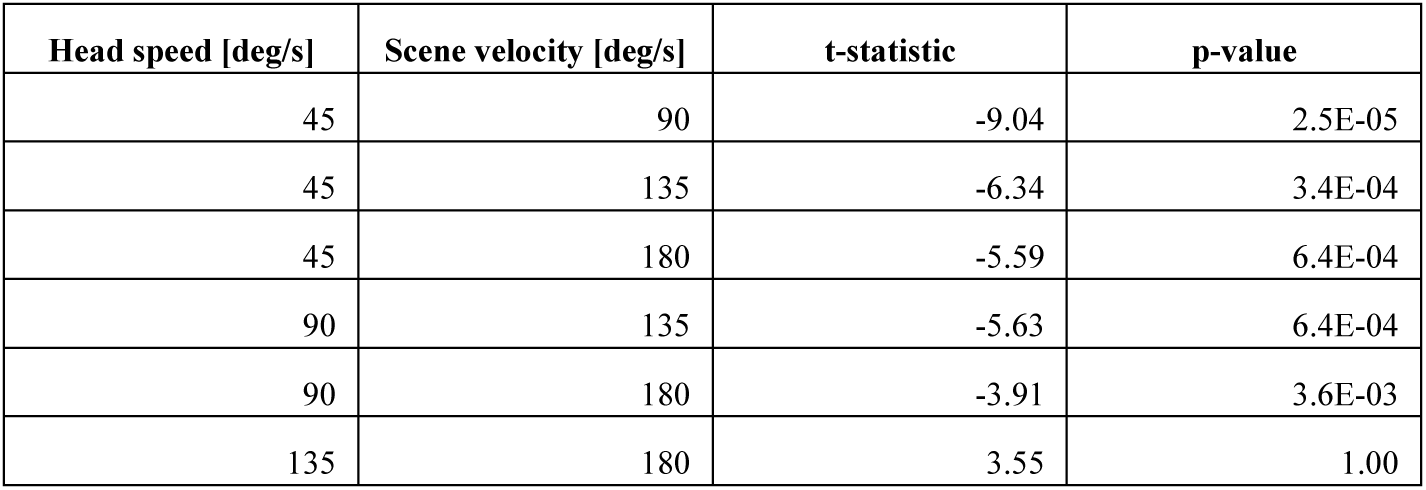

| Head speed [deg/s] | Scene velocity [deg/s] | t-statistic | p-value |
| --- | --- | --- | --- |
| 45 | 90 | -9.04 | 2.5E-05 |
| 45 | 135 | -6.34 | 3.4E-04 |
| 45 | 180 | -5.59 | 6.4E-04 |
| 90 | 135 | -5.63 | 6.4E-04 |
| 90 | 180 | -3.91 | 3.6E-03 |
| 135 | 180 | 3.55 | 1.00 |

We tested the null hypothesis that the animal-averaged decoded head-direction shift evoked by head-only rotations in brightness as well as darkness was the same in wild-type and *FRMD7^tm^*animals. For bright conditions, a one-sided two-sample t-test (n=10 wild-type animals, n=4 *FRMD7^tm^* animals, 12 degrees of freedom) was conducted with the alternative hypothesis that wild-type animals exhibited a greater decoded head-direction shift than *FRMD7^tm^*animals while for dark conditions, the t-test was two-sided. Bonferroni-Holm correction was applied for each bright and dark conditions. The t-statistics and p-values are listed in the table below.

| Head speed [deg/s] | t-statistic, bright | p-value, bright | t-statistic, dark | p-value, dark |
| --- | --- | --- | --- | --- |
| 45 | -8.08 | 5.0E-06 | -1.09 | 1.00 |
| 90 | -7.78 | 5.0E-06 | 1.02 | 1.00 |
| 135 | -9.00 | 2.0E-06 | 1.19 | 1.00 |
| 180 | -5.30 | 9.5E-05 | 0.03 | 1.00 |

A one-sided t-test on related samples (n=10 animals, 9 degrees of freedom) was used to test the null hypothesis that the difference between the angular shifts of decoded and actual head direction is the same for CW and CCW scene rotations (Fig. 5c). The test was performed on the average difference across trials; we repeated both CW and CCW scene rotations 15 times per animal. The t-statistic took the value 6.838, resulting in a p-value of 3.8×10^-5^. An analogous test was performed on *FRMD7^tm^* mice (Fig. 5f; n=4 animals, 3 degrees of freedom), yielding a t-statistic of 0.331 and a p-value of 0.38.

For statistical analysis of escape angles (Fig. 5i,j), we performed two-sided Wilcoxon signed-rank tests on the sine of the difference between the escape angles in the platform-only trial and the control trial, the platform-only trial and the platform-and-scene trial, and the platform-and-scene trial and control trial, respectively. For wild-type mice, the test statistic (W) of the Wilcoxon signed-rank test was 66 for control vs. platform-only, 97 for control vs. platform- and-scene, and 95 for platform-only vs. platform-and-scene. Bonferroni-Holm correction for three comparisons was applied to correct for multiple comparisons. Corrected p-values for the three comparisons were as follows: 0.4263 for control vs. platform-only, 0.0092 for control vs. platform-and-scene, 0.0105 for platform-only vs. platform-and-scene. For *FRMD7^tm^* mice the W statistic was 9 for control vs. platform-only, 10 for control vs. platform-and-scene, and 7 for platform-only vs. platform-and-scene. The Bonferroni-Holm corrected p-values were: 1.875 for control vs. platform-only, 1.875 for control vs. platform-and-scene, 1.625 for platform-only vs. platform-and-scene. The experiment was performed with n=14 wild-type mice and n=5 *FRMD7^tm^* mice, with each mouse receiving one trial of each condition (control, platform-only, platform-and-scene).

### Software

In addition to the software already mentioned, data was analyzed using Python (version 3.11.8)^85^ and the libraries NumPy (1.24.3)^86^, SciPy (1.11.4)^87^, Scikit-learn (1.3.0)^64^, statsmodels (0.14.0)^88^ and pandas (1.5.3)^89^, and visualized using Matplotlib (3.8.0)^90^, seaborn (0.12.2)^91^ and cmocean (3.0.3)^92^.

---

**Extended Data Fig.1.**
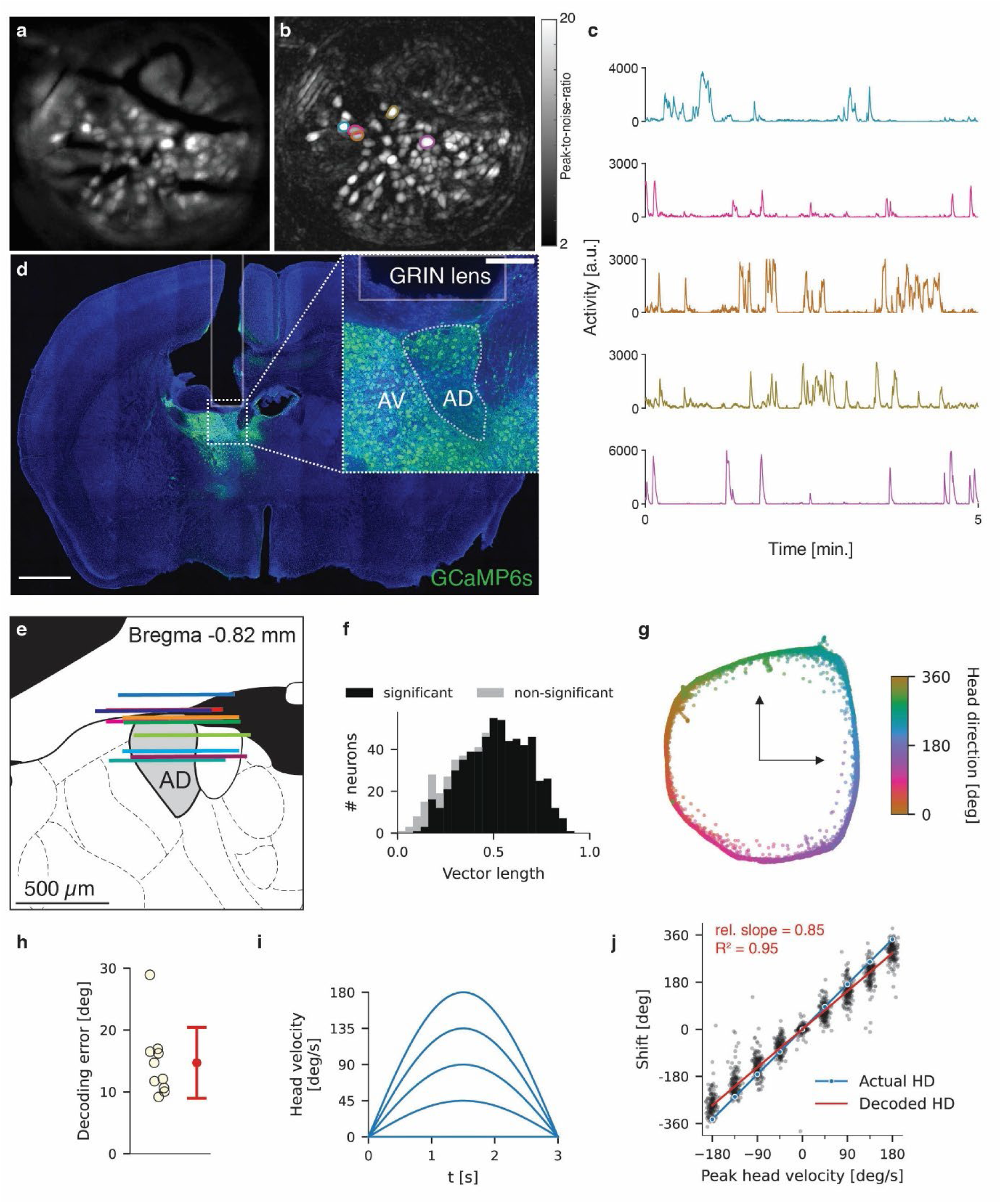
Recording from head-direction cells and decoding head direction from AD population activity. **a**, Maximum intensity projection image from one example recording (same as in Fig. 1b-d,g) in which mice were freely moving for 13 minutes and then passively rotated under head-fixation. **b**, Peak-to-noise-ratio (PNR) image from the same recording as in **a**. For a definition of the PNR, see Ref.^36^. The outline contours of five example cells are shown. **c**, Temporal calcium activity traces for the five example cells outlined in **b** during 5 minutes of free locomotion. **d**, Example coronal brain section showing histological confirmation of GCaMP6s expression and GRIN lens location in the AD. AV: anteroventral nucleus of thalamus. Scale bar: 1 mm. Inset scale bar: 0.2 mm. **e**, Schematic of coronal brain section from a mouse brain atlas^57^ showing the AD and the GRIN lens positions for all 10 wild-type mice (colored lines, each color indicates a different mouse) **f**, Histogram of Rayleigh vector lengths of all cells from all freely moving recordings (n=10 mice). A cell was considered significantly tuned to head direction if its vector length exceeded the 99th percentile of a sample of randomized vector lengths. **g**, Population activity from an example recording during free locomotion (same as **a**) embedded in a two-dimensional plane using Laplacian Eigenmaps. Each data point corresponds to the population activity from a single recording frame and is color-coded according to the actual head direction of the mouse at the corresponding time point. **h**, Decoding error (in degrees) across the entire 13-minute freely moving recording for all mice (n=10). The error bar shows mean ± standard deviation across mice. **i**, Velocity profiles of the platform rotations with peak velocities of 0, 45, 90, 135, 180 degrees per second. **j**, Scatter plot of decoded head-direction shift vs. peak head velocity for all trials and all mice (n=10) in the experiments with passive rotations in the presence of a stationary scene. The blue line shows the angular platform shift as a function of the peak platform velocity, and the red line shows a linear fit of the data (using a Huber regression that is robust to outliers): the slope of the linear fit is 85% of the slope of the curve of the actual head-direction shift, i.e., the decoded head-direction shift underestimates the head shift by 15%.

**Extended Data Fig.2.**
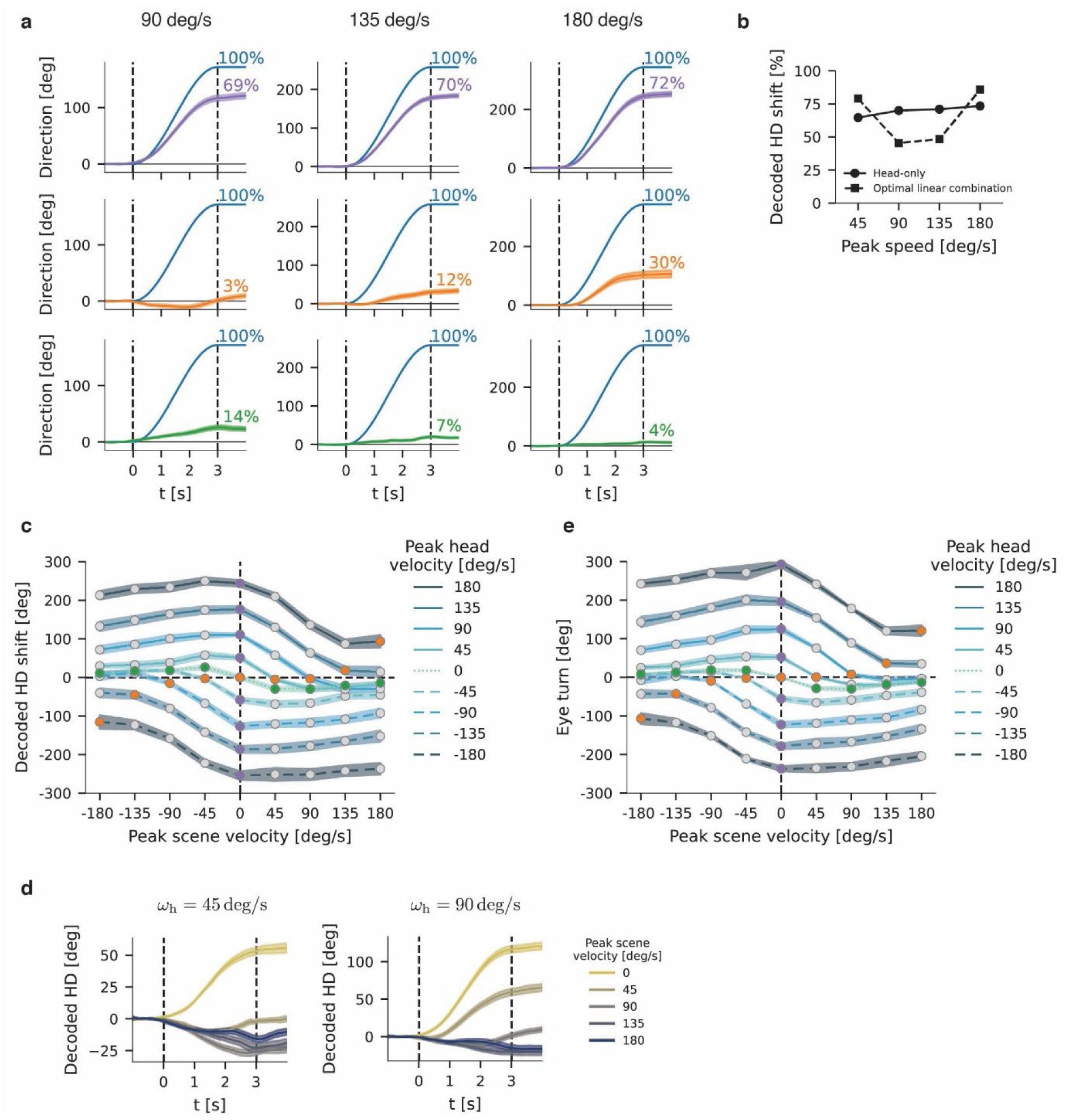
Decoded head-direction and eye turn during visual-vestibular stimulation. **a**, Time series of decoded head direction (purple, orange and green lines) and head and/or scene angle (blue lines) for stimuli with a peak stimulus speed of 90 deg/s (left), 135 deg/s (middle) and 180 deg/s (right). The top row corresponds to head-only rotations, the middle row to head-and-scene rotations and the bottom row to scene-only rotations. Traces show the average across (n=10) animals and the shaded area indicates the standard error of the mean. Note that data from the two experimental conditions involving CW head rotations was sign-inverted to align with CCW head rotations while data from CCW scene rotations was sign-inverted to align with CW scene rotations. **b**, Decoded head-direction shift in head-only rotation (solid line) and optimal linear combination of decoded head-direction shifts in head-and-scene and scene-only rotations (dashed line) as a function of peak stimulus speed. The coefficients of 2.53 on the head-and-scene curve and 2.47 on the scene-only curve were obtained by minimizing the residual sum of squares. **c**, Decoded head-direction shift for the full set of stimulus conditions. Each line shows the angular shift as a function of peak scene velocity for a given value of the peak head speed. Lines are color-coded according to the magnitude of the peak head speed. Solid lines correspond to CW head rotations, dashed lines to CCW head rotations and the dotted line to scene-only rotations. Markers are colored to highlight the three main conditions (purple: head-only rotation, orange: head-and-scene rotation, green: scene-only rotation, grey: remaining conditions) from Fig. 1j. **d**, Time series of decoded head direction for head rotations with a peak speed of 45 deg/s (left) and 90 deg/s (right) for different peak scene velocities (color-coded), where the scene rotates in the same direction as the head. **e**, Eye turn in degrees for the full set of experimental conditions, analogous to **c**, for (n=7) animals in which the eye was tracked.

**Extended Data Fig.3.**
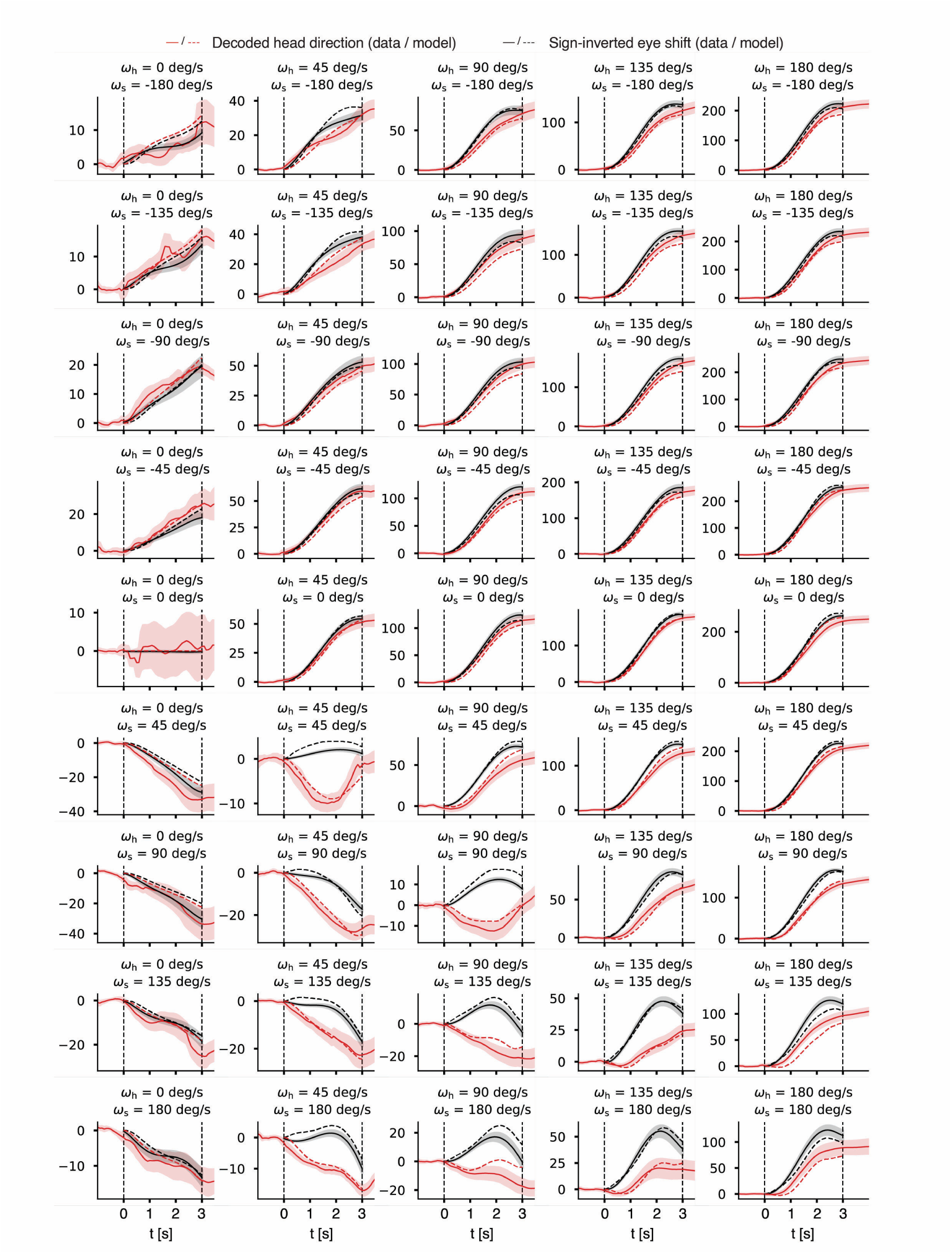
Time series of decoded head direction and sign-inverted eye shift for each visual-vestibular stimulation condition. Time series of the decoded head direction (red lines), the sign-inverted angular eye shift (black lines) for each experimental condition corresponding to different combinations of peak head velocity (ω_h_) and peak scene velocity (ω_s_). Solid lines correspond to the data and dashed lines correspond to model predictions. Data traces show the average across animals (n=7 mice) after aggregation of data from CW and CCW head rotations. The shaded area indicates the standard error of the mean. The y-axis is in degrees.

**Extended Data Fig.4.**
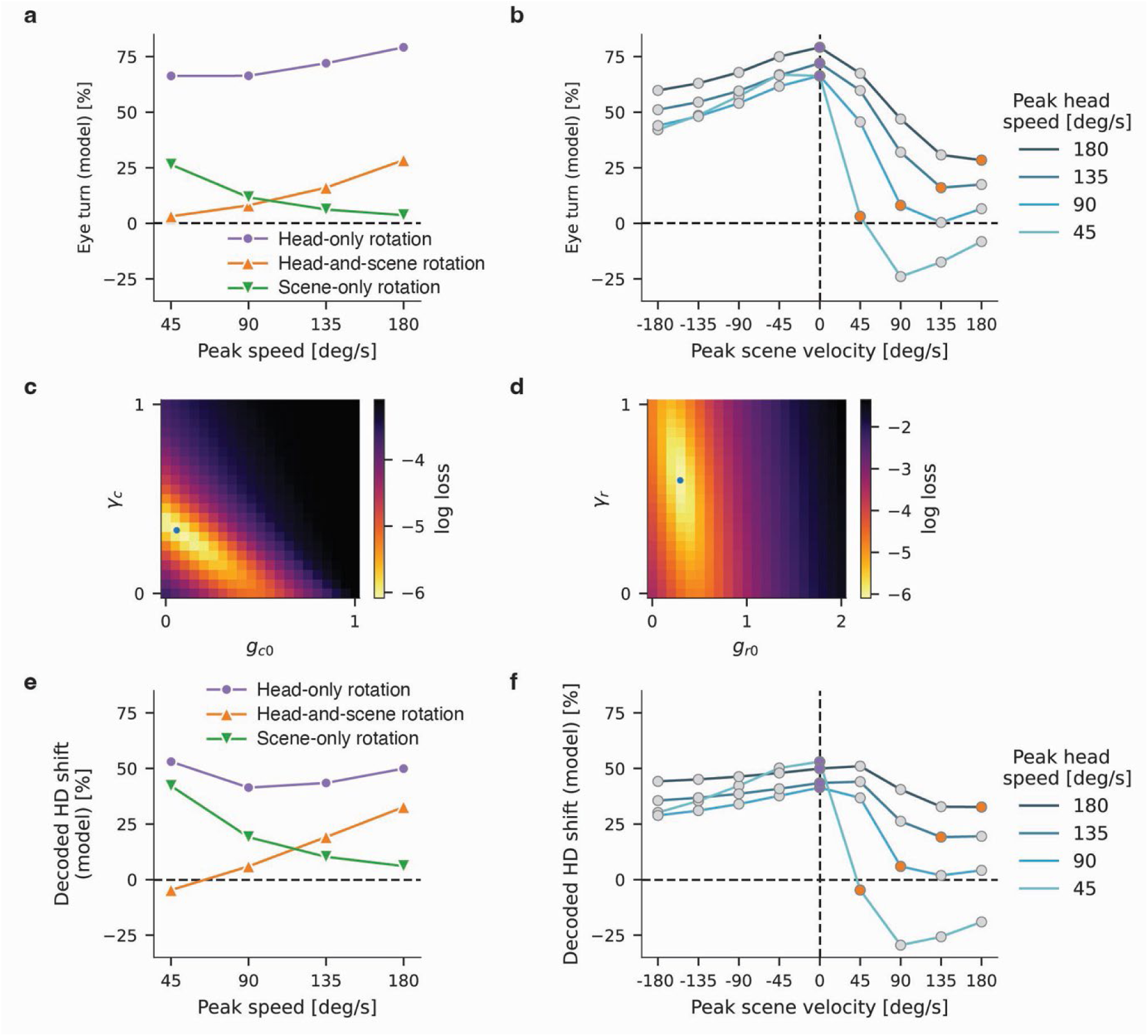
Model predictions of eye turn and model sensitivity analyses. **a**-**b**, Model prediction (displayed as a percentage of the stimulus rotation) for the eye turn. **a**, Model prediction as a function of peak stimulus speed for head-only (purple line), head-and-scene (orange line) and scene-only rotation (green line). **b**, Model prediction as a function of peak scene velocity for each value of the peak head speed (color-coding). **c**, Heat map of the logarithmic loss corresponding to the error between the model prediction and the data (see Methods) dependent on the vestibular gain parameters. The gain *g*_*c*_ on the canal signal *c* is given by *g*_*c*_ = *g*_*c*0_ + χ_*c*_ |*c*|. **d**, Sensitivity analysis of the loss as in **c** but for the parameters of the gain on the visual signal; the gain is given by *g*_*r*_ = *g*_*r*0_ + χ_*r*_|*c*|. **e**-**f**, Model prediction for the decoded head-direction shift (displayed as a percentage of angular stimulus shift) when the model is fit with a constant gain on the retinal slip, i.e. with χ_*r*_ = 0. **e**, Model prediction as a function of peak stimulus speed for head-only rotations (purple line), head-and-scene rotations (orange line) and scene-only rotations (green line). **f**, Model prediction as a function of peak scene velocity for each value of the peak head speed (color-coding).

**Extended Data Fig.5.**
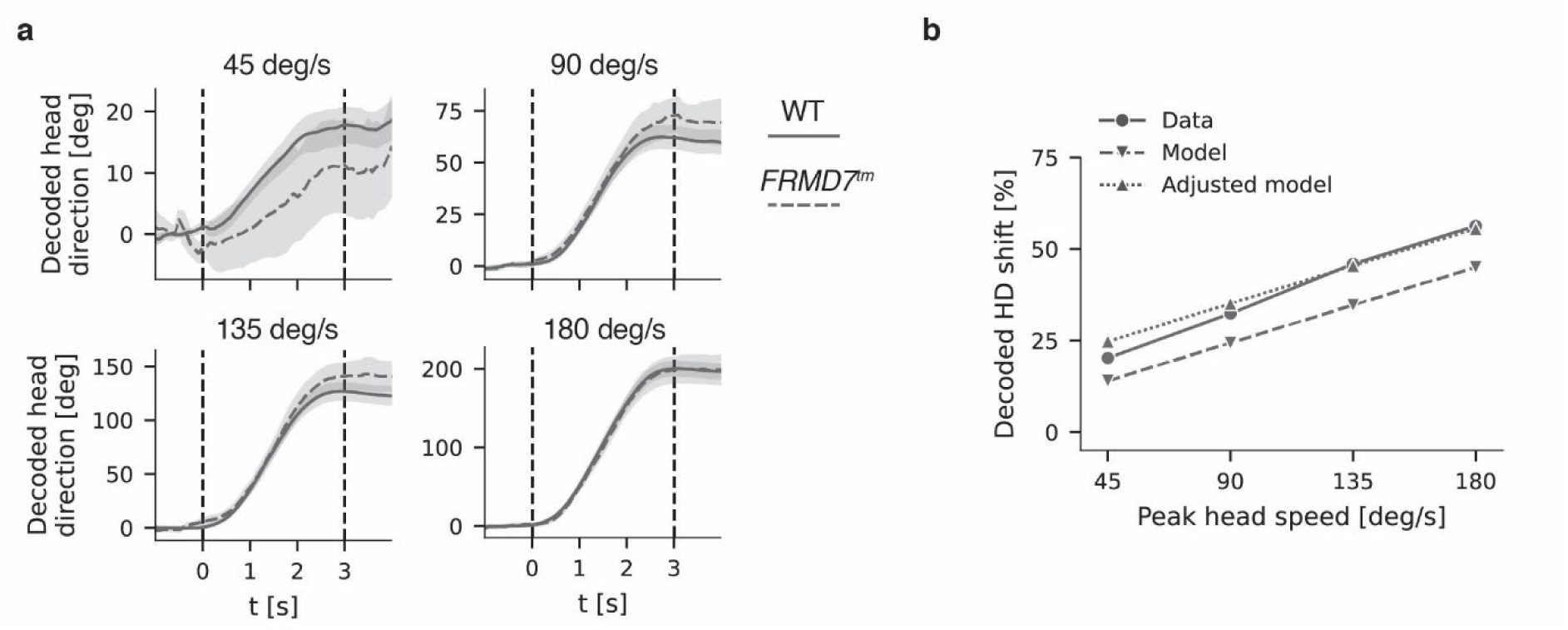
Time series of decoded head direction in darkness and model prediction of decoded head-direction shift with adjusted vestibular gain. **a**, Time series of decoded head direction for wild-type (solid line) and *FRMD7^tm^* mice (dashed line) for head rotations in darkness with peak head speed of 45, 90, 135 or 180 deg/s. Lines correspond to the average across (n=10 wild-type respectively n=4 *FRMD7^tm^*) mice and the shaded areas indicate the standard error of the mean. **b**, Decoded head-direction shift in percent of the angular head shift as a function of the peak head speed for wild-type mice (solid line, average across the same n=7 wild-type mice used for fitting the other model parameters) and the model prediction with (dotted line) and without (dashed line) adjustment of the parameter *g*_*c*0_ of the gain on the canal signal.

**Extended Data Fig.6.**
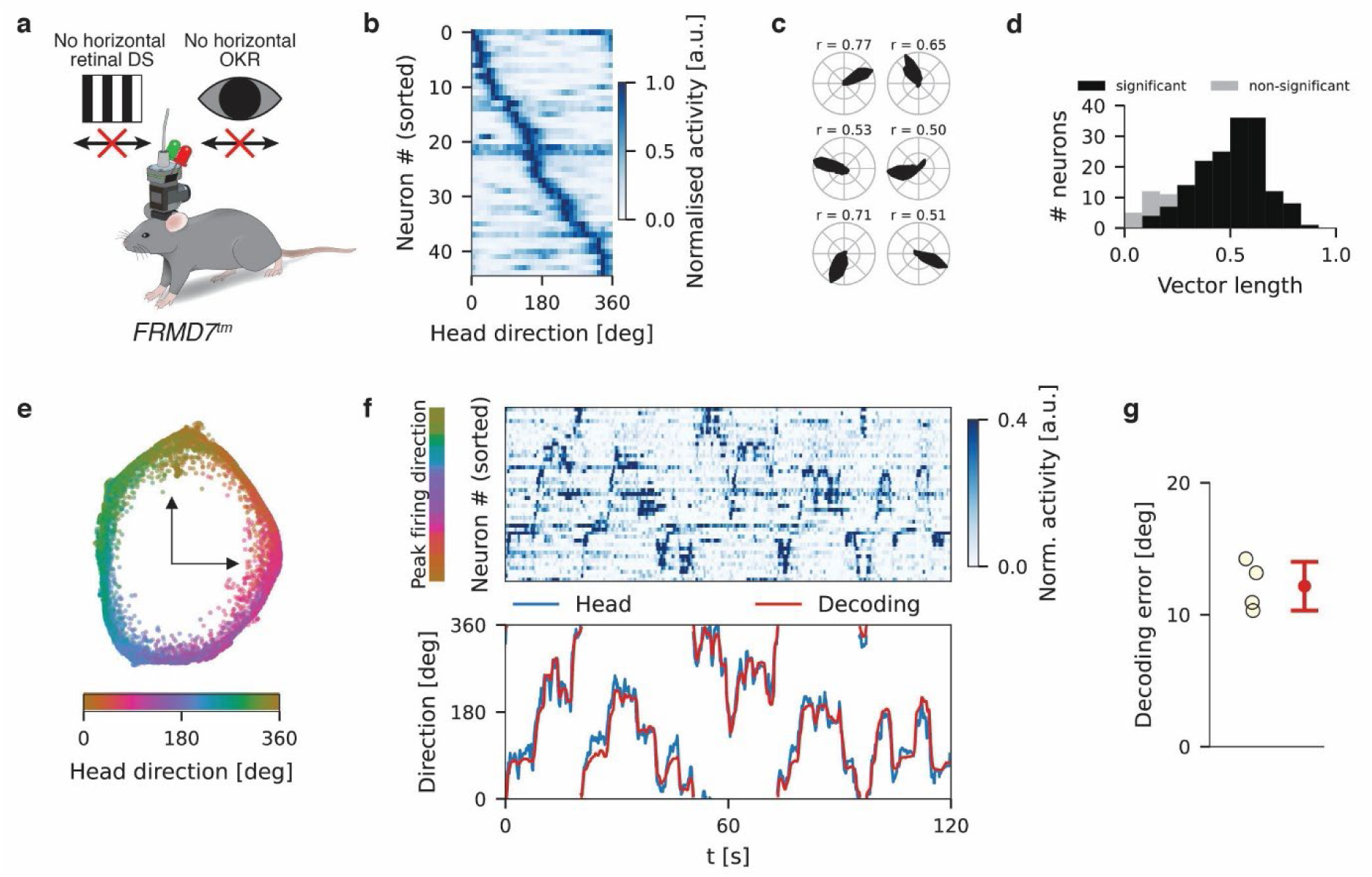
Head-direction tuning is intact in freely moving *FRMD7^tm^* mice. **a**, Schematic of recording from *FRMD7^tm^* mice with the miniature head-mounted microscope. **b**, Head-direction tuning of all extracted cells from one example recording during free locomotion. **c**, Polar plots and Rayleigh vector lengths of example cells from the same recording as in **b**. **d**, Histogram of Rayleigh vector lengths of all cells from all recordings (n=4 mice). A cell was considered significantly tuned to head direction if its vector length exceeded the 99th percentile of a sample of randomized vector lengths. **e**, Embedding of the population activity in a two-dimensional plane using Laplacian Eigenmaps, from the same recording as in **b**. Each data point corresponds to the population activity from a single recording frame and is color-coded according to the actual head direction of the mouse at the corresponding time point. **f**, Top: Normalized activity of all extracted cells (sorted by peak firing direction) during a 30 second time window from the same recording as in **b**. Bottom: Actual (blue) and decoded (red) head direction for the same time window shown above. **g**, Decoding error (in degrees) across the entire freely moving recording for all mice (n=4).

**Extended Data Fig.7.**
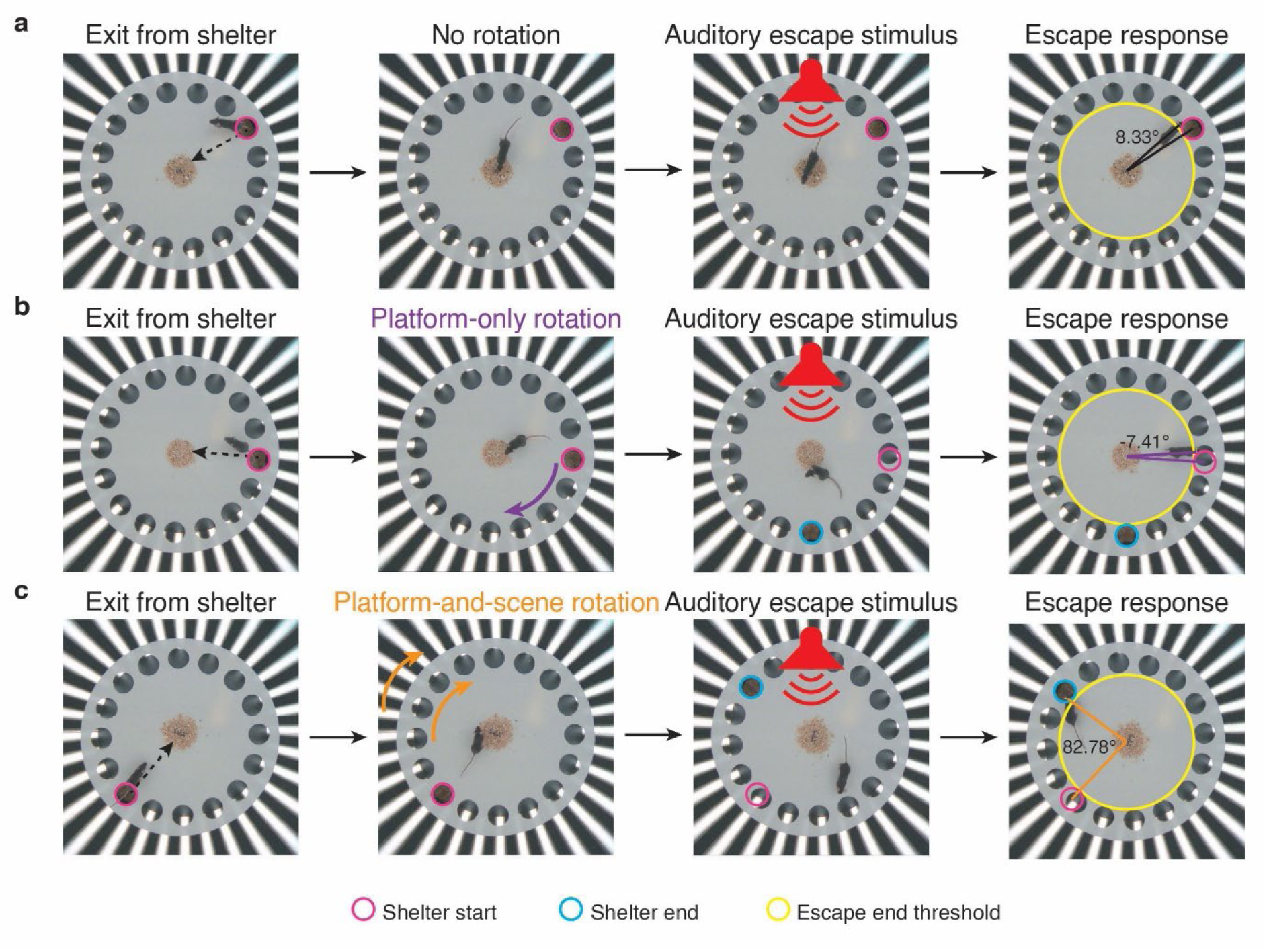
Calculation of escape endpoint angle in the escape-to-shelter behavior paradigm. **a**, Frames from the behavioral video showing the sequence of events during a control trial, in which neither the platform nor the scene was rotated. From left to right: i) The mouse exits the shelter and walks towards the center of the platform. ii) The mouse reaches the center of the platform. iii) An auditory stimulus is presented that triggers an escape response. iv) The end of the escape response: defined as the moment when both ears of the mouse cross the circular boundary touching the potential shelter locations (indicated by the yellow circle). The angle shows the escape endpoint angle. The escape endpoint angle corresponds to the angle between the vector from the platform center to the head of the mouse (the center of a line connecting the two ears) and the vector from the platform center to the center of the shelter. The angles clockwise to the shelter were negative and angles counterclockwise to the shelter were positive. **b**, Same as **a** for a platform-only rotation trial. From left to right: i) The mouse exits the shelter and walks towards the center of the platform. ii) The mouse reaches the center of the platform and the platform rotation begins iii) Immediately after the platform rotation is complete the auditory stimulus is presented. iv) The end of the escape response: defined as the angle between the vector from the platform center to the head of the mouse and the vector from the platform center to the center of the shelter starting position, at the time when both ears of the mouse crossed the boundary indicated by the yellow circle. Escape endpoint angles from trials with clockwise platform rotation were multiplied by -1 to align data from clockwise and counterclockwise platform rotations. Therefore, an escape to the shelter end position always equaled 85.9 degrees regardless of the direction of the platform rotation. **c**, Same as **b**, except both the platform and scene were rotated in synchrony.

**Extended Data Fig.8.**
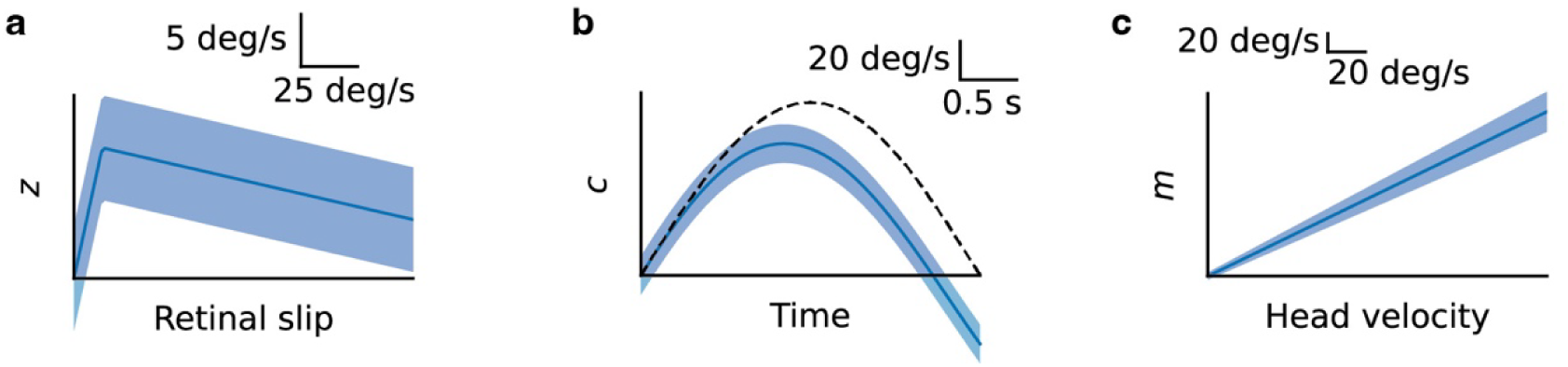
Signal and noise structure assumed in the Bayesian model. **a**, The solid line corresponds to the mean visual signal as a function of retinal slip, the shaded area indicates the standard deviation. **b**, The solid line corresponds to the vestibular signal over time for the experimentally applied head rotation stimulus with a peak head velocity of 90 deg/s, obtained from a model of sensory transduction in the semicircular canals. The shaded area indicates the standard deviation of the vestibular signal. Note that, for simplicity, the Bayesian model assumes that the vestibular signal matches the true head velocity on average. **c**, The solid line corresponds to the mean motor signal and the shaded area indicates the standard deviation as a function of head velocity.

**Extended Data Fig.9.**
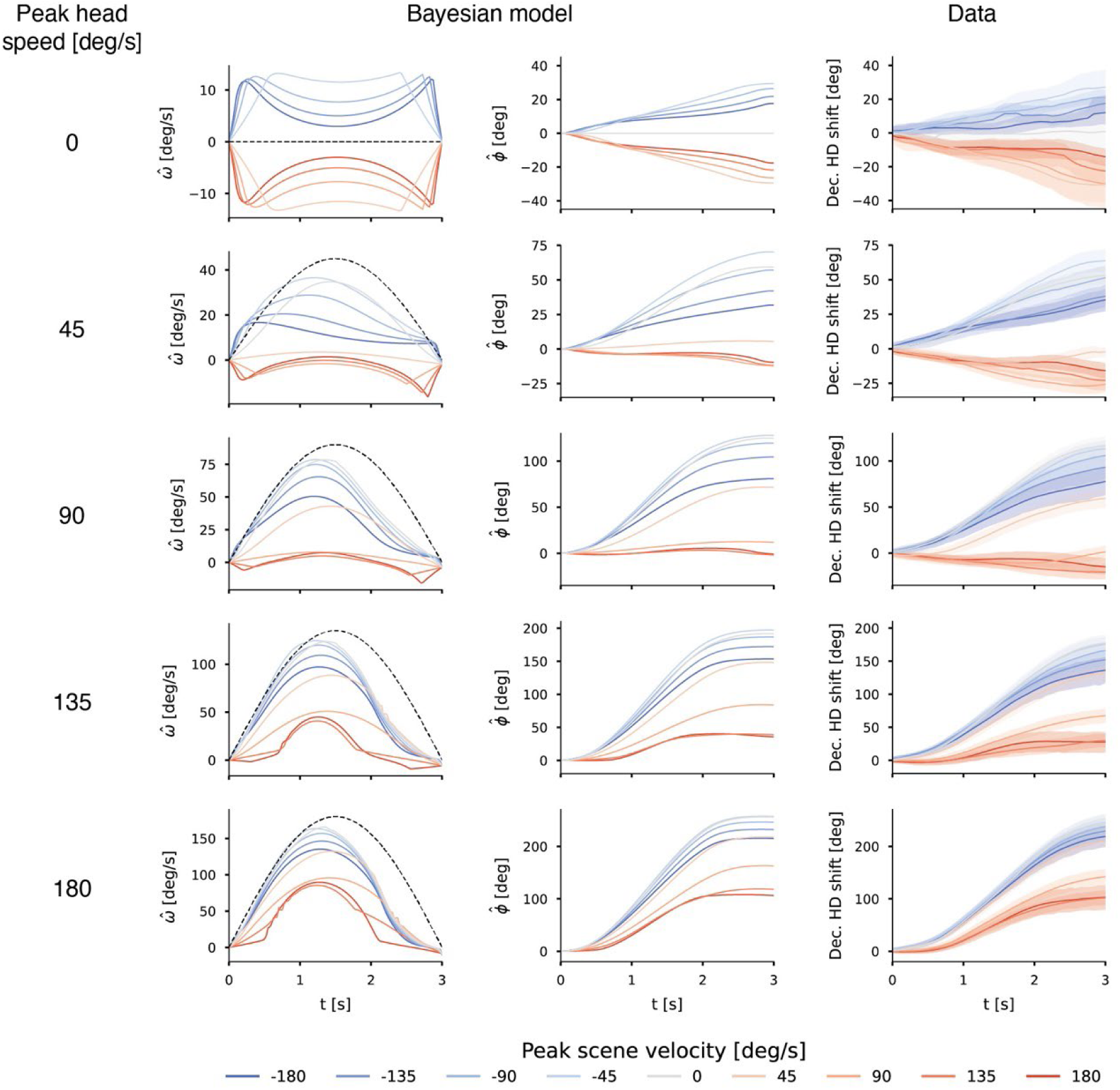
Dynamics of Bayesian estimation for the experimentally applied rotation stimuli and comparison to measured decoded head-direction shift. Left: Time series of the posterior mean head velocity. Center: Cumulative integral of the posterior mean head velocity over time, representing estimated head-direction shift. Right: Time series of the measured decoded head-direction shift (solid lines: mean, shaded areas: standard error of the mean). Each row corresponds to a different peak head speed (indicated on the left). Colors denote different peak scene velocities as shown in the legend below.

**Supplementary Video 1.** Examples of scene rotation trials during calcium imaging from the AD in a freely moving mouse.

**Supplementary Video 2.** An example of a control trial from the spatial navigation paradigm. Note that audio was not recorded by the camera; the auditory stimulus used in the experiment (an owl vocalization) was edited into the video file for illustrative purposes.

**Supplementary Video 3.** An example of a platform-only rotation trial from the spatial navigation paradigm (same mouse as in Supplementary Video 2). Note that audio was not recorded by the camera; the auditory stimulus used in the experiment (an eagle vocalization) was edited into the video file for illustrative purposes.

**Supplementary Video 4.** An example of a platform-and-scene rotation trial from the spatial navigation paradigm (same mouse as in Supplementary Video 2). Note that audio was not recorded by the camera; the auditory stimulus used in the experiment (a hawk vocalization) was edited into the video file for illustrative purposes.

## References

1. Tolman, E. C. Cognitive maps in rats and men. Psychol. Rev. 55, 189–208 (1948).

2. Behrens, T. E. J. et al. What Is a Cognitive Map? Organizing Knowledge for Flexible Behavior. Neuron 100, 490–509 (2018).

3. O’Keefe, J. Place units in the hippocampus of the freely moving rat. Exp. Neurol. 51, 78–109 (1976).

4. O’Keefe, J. & Nadel, L. The Hippocampus as a Cognitive Map. (Oxford University Press, 1978).

5. Hafting, T., Fyhn, M., Molden, S., Moser, M.-B. & Moser, E. I. Microstructure of a spatial map in the entorhinal cortex. Nature 436, 801–806 (2005).

6. Taube, J., Muller, R. & Ranck, J. Head-direction cells recorded from the postsubiculum in freely moving rats. I. Description and quantitative analysis. J. Neurosci. 10, 420–435 (1990).

7. Taube, J., Muller, R. & Ranck, J. Head-direction cells recorded from the postsubiculum in freely moving rats. II. Effects of environmental manipulations. J. Neurosci. 10, 436–447 (1990).

8. Etienne, A. S. & Jeffery, K. J. Path integration in mammals. Hippocampus 14, 180–192 (2004).

9. Valerio, S. & Taube, J. S. Path integration: how the head direction signal maintains and corrects spatial orientation. Nat. Neurosci. 15, 1445–1453 (2012).

10. Butler, W. N., Smith, K. S., Van Der Meer, M. A. A. & Taube, J. S. The Head-Direction Signal Plays a Functional Role as a Neural Compass during Navigation. Curr. Biol. 27, 1259–1267 (2017).

11. Cullen, K. E. & Taube, J. S. Our sense of direction: progress, controversies and challenges. Nat. Neurosci. 20, 1465–1473 (2017).

12. Peyrache, A., Lacroix, M. M., Petersen, P. & Buzsáki, G. Internally-organized mechanisms of the head direction sense. Nat. Neurosci. 18, 569–575 (2015).

13. Robertson, R. G., Rolls, E. T., Georges-François, P. & Panzeri, S. Head direction cells in the primate pre-subiculum. Hippocampus 9, 206–219 (1999).

14. Muir, G. M. et al. Disruption of the head direction cell signal after occlusion of the semicircular canals in the freely moving chinchilla. J. Neurosci. Off. J. Soc. Neurosci. 29, 14521–14533 (2009).

15. Seelig, J. D. & Jayaraman, V. Neural dynamics for landmark orientation and angular path integration. Nature 521, 186–191 (2015).

16. Finkelstein, A. et al. Three-dimensional head-direction coding in the bat brain. Nature 517, 159–164 (2015).

17. Varga, A. G. & Ritzmann, R. E. Cellular Basis of Head Direction and Contextual Cues in the Insect Brain. Curr. Biol. 26, 1816–1828 (2016).

18. Petrucco, L. et al. Neural dynamics and architecture of the heading direction circuit in zebrafish. Nat. Neurosci. 26, 765–773 (2023).

19. Ben-Yishay, E. et al. Directional tuning in the hippocampal formation of birds. Curr. Biol. 31, 2592–2602.e4 (2021).

20. Skaggs, W. E., Knierim, J. J., Kudrimoti, H. S. & McNaughton, B. L. A model of the neural basis of the rat’s sense of direction. Adv. Neural Inf. Process. Syst. 7, 173–180 (1995).

21. Zhang, K. Representation of spatial orientation by the intrinsic dynamics of the head-direction cell ensemble: a theory. J. Neurosci. 16, 2112–2126 (1996).

22. Goodridge, J. P., Dudchenko, P. A., Worboys, K. A., Golob, E. J. & Taube, J. S. Cue control and head direction cells. Behav. Neurosci. 112, 749–761 (1998).

23. Zugaro, M. B., Tabuchi, E. & Wiener, S. I. Influence of conflicting visual, inertial and substratal cues on head direction cell activity. Exp. Brain Res. 133, 198–208 (2000).

24. Yoder, R. M., Clark, B. J. & Taube, J. S. Origins of landmark encoding in the brain. Trends Neurosci. 34, 561–571 (2011).

25. Stackman, R. W. & Taube, J. S. Firing Properties of Head Direction Cells in the Rat Anterior Thalamic Nucleus: Dependence on Vestibular Input. J. Neurosci. 17, 4349–4358 (1997).

26. Stackman, R. W., Clark, A. S. & Taube, J. S. Hippocampal spatial representations require vestibular input. Hippocampus 12, 291–303 (2002).

27. Blair, H. T. & Sharp, P. E. Visual and vestibular influences on head-direction cells in the anterior thalamus of the rat. Behav. Neurosci. 110, 643–660 (1996).

28. Arleo, A. et al. Optic Flow Stimuli Update Anterodorsal Thalamus Head Direction Neuronal Activity in Rats. J. Neurosci. 33, 16790–16795 (2013).

29. Ajabi, Z., Keinath, A. T., Wei, X.-X. & Brandon, M. P. Population dynamics of head-direction neurons during drift and reorientation. Nature 615, 892–899 (2023).

30. Leigh, R. J. & Zee, D. S. The Neurology of Eye Movements. (Oxford University Press, Oxford, 2015).

31. Schweigart, G., Mergner, T., Evdokimidis, I., Morand, S. & Becker, W. Gaze Stabilization by Optokinetic Reflex (OKR) and Vestibulo-ocular Reflex (VOR) During Active Head Rotation in Man. Vision Res. 37, 1643–1652 (1997).

32. Taube, J. S. The Head Direction Signal: Origins and Sensory-Motor Integration. Annu. Rev. Neurosci. 30, 181–207 (2007).

33. Yonehara, K. et al. Congenital Nystagmus Gene FRMD7 Is Necessary for Establishing a Neuronal Circuit Asymmetry for Direction Selectivity. Neuron 89, 177–193 (2016).

34. Belkin, M. & Niyogi, P. Laplacian Eigenmaps for Dimensionality Reduction and Data Representation. Neural Comput. 15, 1373–1396 (2003).

35. Shinder, M. E. & Taube, J. S. Active and passive movement are encoded equally by head direction cells in the anterodorsal thalamus. J. Neurophysiol. 106, 788–800 (2011).

36. Zhou, P. et al. Efficient and accurate extraction of in vivo calcium signals from microendoscopic video data. eLife 7, e28728 (2018).

37. Giovannucci, A. et al. CaImAn an open source tool for scalable calcium imaging data analysis. eLife 8, e38173 (2019).

38. Van Alphen, A. M., Stahl, J. S. & De Zeeuw, C. I. The dynamic characteristics of the mouse horizontal vestibulo-ocular and optokinetic response. Brain Res. 890, 296–305 (2001).

39. Iwashita, M., Kanai, R., Funabiki, K., Matsuda, K. & Hirano, T. Dynamic properties, interactions and adaptive modifications of vestibulo-ocular reflex and optokinetic response in mice. Neurosci. Res. 39, 299– 311 (2001).

40. Lasker, D. M., Han, G. C., Park, H. J. & Minor, L. B. Rotational Responses of Vestibular–Nerve Afferents Innervating the Semicircular Canals in the C57BL/6 Mouse. J. Assoc. Res. Otolaryngol. 9, 334–348 (2008).

41. Dhande, O. S. et al. Genetic Dissection of Retinal Inputs to Brainstem Nuclei Controlling Image Stabilization. J. Neurosci. 33, 17797–17813 (2013).

42. Robinson, D. A. Models of the optokinetic system. in Progress in Brain Research vol. 267 231–249 (Elsevier, 2022).

43. Schmid, R., Buizza, A. & Zambarbieri, D. A non-linear model for visual-vestibular interaction during body rotation in man. Biol. Cybern. 36, 143–151 (1980).

44. Stahl, J. S. & Thumser, Z. C. Dynamics of abducens nucleus neurons in the awake mouse. J. Neurophysiol. 108, 2509–2523 (2012).

45. Stahl, J. S. et al. Mechanics of mouse ocular motor plant quantified by optogenetic techniques. J. Neurophysiol. 114, 1455–1467 (2015).

46. Yoshida, K. et al. A Key Role of Starburst Amacrine Cells in Originating Retinal Directional Selectivity and Optokinetic Eye Movement. Neuron 30, 771–780 (2001).

47. Mongeau, R., Miller, G. A., Chiang, E. & Anderson, D. J. Neural Correlates of Competing Fear Behaviors Evoked by an Innately Aversive Stimulus. J. Neurosci. 23, 3855–3868 (2003).

48. Vale, R., Evans, D. A. & Branco, T. Rapid Spatial Learning Controls Instinctive Defensive Behavior in Mice. Curr. Biol. 27, 1342–1349 (2017).

49. Meyer, A. F., O’Keefe, J. & Poort, J. Two Distinct Types of Eye-Head Coupling in Freely Moving Mice. Curr. Biol. 30, 2116–2130.e6 (2020).

50. Michaiel, A. M., Abe, E. T. & Niell, C. M. Dynamics of gaze control during prey capture in freely moving mice. eLife 9, e57458 (2020).

51. Senzai, Y. & Scanziani, M. A cognitive process occurring during sleep is revealed by rapid eye movements. Science 377, 999–1004 (2022).

52. Land, M. F. Eye movements of vertebrates and their relation to eye form and function. J. Comp. Physiol. A 201, 195–214 (2015).

53. Hulse, B. K. & Jayaraman, V. Mechanisms Underlying the Neural Computation of Head Direction. Annu. Rev. Neurosci. 43, 31–54 (2020).

54. Peirce, J. et al. PsychoPy2: Experiments in behavior made easy. Behav. Res. Methods 51, 195–203 (2019).

55. Schindelin, J., et al. Fiji: an open-source platform for biological-image analysis. Nat. Methods 9, 676–682 (2012).

56. Berens, P. CircStat: A MATLAB Toolbox for Circular Statistics. J. Stat. Softw. 31, 1–21 (2009).

57. Paxinos, G. & Franklin, K. B. J. The Mouse Brain in Stereotaxic Coordinates: Compact Second Edition. (Gulf Professional Publishing, 2004).

58. Geng, J. et al. Chronic Ca2+ imaging of cortical neurons with long-term expression of GCaMP-X. eLife 11, e76691 (2022).

59. Friedrich, J., Zhou, P. & Paninski, L. Fast online deconvolution of calcium imaging data. PLOS Comput. Biol. 13, e1005423 (2017).

60. Chen, T.-W. et al. Ultrasensitive fluorescent proteins for imaging neuronal activity. Nature 499, 295–300 (2013).

61. Bradski, G. The opencv library. Dr Dobb’s J. Softw. Tools 25, 120–123 (2000).

62. Moritz, P. et al. Ray: A Distributed Framework for Emerging AI Applications. in Proceedings of the 13th USENIX Symposium on Operating Systems Design and Implementation (OSDI ’18) 561–577 (2018).

63. Coifman, R. R. & Lafon, S. Diffusion maps. Appl. Comput. Harmon. Anal. 21, 5–30 (2006).

64. Pedregosa, F. et al. Scikit-learn: Machine Learning in Python. J. Mach. Learn. Res. 12, 2825–2830 (2011).

65. Fisher, N. I. & Lee, A. J. A correlation coefficient for circular data. Biometrika 70, 327–332 (1983).

66. Blair, H. & Sharp, P. Anticipatory head direction signals in anterior thalamus: evidence for a thalamocortical circuit that integrates angular head motion to compute head direction. J. Neurosci. 15, 6260–6270 (1995).

67. Blair, H. T., Lipscomb, B. W. & Sharp, P. E. Anticipatory Time Intervals of Head-Direction Cells in the Anterior Thalamus of the Rat: Implications for Path Integration in the Head-Direction Circuit. J. Neurophysiol. 78, 145–159 (1997).

68. Blair, H. T., Cho, J. & Sharp, P. E. Role of the Lateral Mammillary Nucleus in the Rat Head Direction Circuit: A Combined Single Unit Recording and Lesion Study. Neuron 21, 1387–1397 (1998).

69. Taube, J. S. & Muller, R. U. Comparisons of head direction cell activity in the postsubiculum and anterior thalamus of freely moving rats. Hippocampus 8, 87–108 (1998).

70. Bassett, J. P. et al. Passive Movements of the Head Do Not Abolish Anticipatory Firing Properties of Head Direction Cells. J. Neurophysiol. 93, 1304–1316 (2005).

71. Mathis, A. et al. DeepLabCut: markerless pose estimation of user-defined body parts with deep learning. Nat. Neurosci. 21, 1281–1289 (2018).

72. Sakatani, T. & Isa, T. PC-based high-speed video-oculography for measuring rapid eye movements in mice. Neurosci. Res. 49, 123–131 (2004).

73. Collewijn, H. An analog model of the rabbit’s optokinetic system. Brain Res. 36, 71–88 (1972).

74. Buizza, A. & Schmid, R. Visual-vestibular interaction in the control of eye movement: Mathematical modelling and computer simulation. Biol. Cybern. 43, 209–223 (1982).

75. Maioli, C. Optokinetic nystagmus: modeling the velocity storage mechanism. J. Neurosci. 8, 821–832 (1988).

76. Cullen, K. E. The neural encoding of self-motion. Curr. Opin. Neurobiol. 21, 587–595 (2011).

77. Fernandez, C. & Goldberg, J. M. Physiology of peripheral neurons innervating semicircular canals of the squirrel monkey. II. Response to sinusoidal stimulation and dynamics of peripheral vestibular system. J. Neurophysiol. 34, 661–675 (1971).

78. Xie, X., Hahnloser, R. H. R. & Seung, H. S. Double-ring network model of the head-direction system. Phys. Rev. E 66, 041902 (2002).

79. Boucheny, C., Brunel, N. & Arleo, A. A continuous attractor network model without recurrent excitation: maintenance and integration in the head direction cell system. J. Comput. Neurosci. 18, 205–227 (2005).

80. Stackman, R. W. & Taube, J. S. Firing Properties of Rat Lateral Mammillary Single Units: Head Direction, Head Pitch, and Angular Head Velocity. J. Neurosci. 18, 9020–9037 (1998).

81. Robinson, D. A. The behavior of motoneurons. in Progress in Brain Research vol. 267 15–42 (Elsevier, 2022).

82. Fitts, P. M. The information capacity of the human motor system in controlling the amplitude of movement. J. Exp. Psychol. 47, 381–391 (1954).

83. Darling, W. G. & Cooke, J. D. Changes in the variability of movement trajectories with practice. J. Mot. Behav. 19, 291–309 (1987).

84. van Beers, R. J., Haggard, P. & Wolpert, D. M. The Role of Execution Noise in Movement Variability. J. Neurophysiol. 91, 1050–1063 (2004).

85. Rossum, G. van & Drake, F. L. The Python Language Reference. (Python Software Foundation, 2010).

86. Harris, C. R. et al. Array programming with NumPy. Nature 585, 357–362 (2020).

87. Virtanen, P. et al. SciPy 1.0: fundamental algorithms for scientific computing in Python. Nat. Methods 17, 261–272 (2020).

88. Seabold, S. & Perktold, J. Statsmodels: Econometric and Statistical Modeling with Python. in PROC. OF THE 9th PYTHON IN SCIENCE CONF 92–96 (SCIPY, 2010).

89. McKinney, W. Data Structures for Statistical Computing in Python. in PROC. OF THE 9th PYTHON IN SCIENCE CONF 56–61 (SCIPY, 2010).

90. Hunter, J. D. Matplotlib: A 2D Graphics Environment. Comput. Sci. Eng. 9, 90–95 (2007).

91. Waskom, M. L. seaborn: statistical data visualization. J. Open Source Softw. 6, 3021 (2021).

92. Thyng, K. M., Greene, C. A., Hetland, R. D., Zimmerle, H. M. & DiMarco, S. F. True Colors of Oceanography: Guidelines for Effective and Accurate Colormap Selection. Oceanography 29, 9–13 (2016).

